# Integrative *in situ* mapping of single-cell transcriptional states and tissue histopathology in an Alzheimer’s disease model

**DOI:** 10.1101/2022.01.14.476072

**Authors:** Hu Zeng, Jiahao Huang, Haowen Zhou, William J. Meilandt, Borislav Dejanovic, Yiming Zhou, Christopher J. Bohlen, Seung-Hye Lee, Jingyi Ren, Albert Liu, Hao Sheng, Jia Liu, Morgan Sheng, Xiao Wang

## Abstract

Amyloid-β plaques and neurofibrillary tau tangles are the neuropathologic hallmarks of Alzheimer’s disease (AD), but the spatiotemporal cellular responses and molecular mechanisms underlying AD pathophysiology remain poorly understood. Here we introduce STARmap PLUS to simultaneously map single-cell transcriptional states and disease marker proteins in brain tissues of AD mouse models at subcellular resolution (200 nm). This high-resolution spatial transcriptomics map revealed a core-shell structure where disease-associated microglia (DAM) closely contact amyloid-β plaques, whereas disease-associated astrocytes (DAA) and oligodendrocyte precursor cells (OPC) are enriched in the outer shells surrounding the plaque- DAM complex. Hyperphosphorylated tau emerged mainly in excitatory neurons in the CA1 region accompanied by the infiltration of oligodendrocyte subtypes into the axon bundles of hippocampal alveus. The integrative STARmap PLUS method bridges single-cell gene expression profiles with tissue histopathology at subcellular resolution, providing an unprecedented roadmap to pinpoint the molecular and cellular mechanisms of AD pathology and neurodegeneration.

## Introduction

Alzheimer’s disease (AD) is a progressive neurodegenerative disorder and the most common cause of dementia in the elderly^1^. Widespread deposition of extracellular amyloid-β (Aβ) plaques and intracellular neurofibrillary tangles (hyperphosphorylated tau deposits), especially in the neocortex and hippocampus, are the neuropathologic hallmarks of AD^1–4^. In addition, AD pathology features reactive changes of microglia and astrocytes and white matter abnormalities^5–7^. A key question in AD research is how the histopathological hallmarks are correlated with molecular disturbance that drives neurodegeneration across different cell populations.

Conventional experimental approaches are disadvantageous for uncovering the molecular and cellular complexity of AD: Bulk-tissue analyses mask the heterogeneity of cell populations in the brain; standard imaging methods visualize only a few genes/proteins and access limited cell types and brain regions. The recent application of single-cell RNA sequencing (scRNA-seq) to AD brain tissue has been transformative, revealing substantial and heterogeneous changes of gene expression in major brain cell types in patients and mouse models^8–13^. A sub-population of microglia with a distinctive transcriptomic state (termed disease-associated microglia or DAM) was identified by bulk and scRNA-seq studies of mouse AD models and other neurodegenerative disorders^10, 14–19^. Besides DAM microglia, disease-associated astrocytes (DAA) with characteristic gene signatures emerge in AD, suggesting a major transcriptional response in AD by multiple cell types^9, 20–25^.

However, there are major limitations of scRNA-seq methods: they cannot preserve the spatial pattern of single cells (or their relationships to localized tissue pathology); and the isolation of single cells or single nuclei from the brain tissue can introduce significant bias in cellular representation and artifactual changes in gene expression^26, 27^. Therefore, a fundamentally different technology platform capable of integrating spatially resolved single-cell transcriptomics with histology and immunostaining in intact tissue is needed to fully understand the scope and heterogeneity of diverse cellular responses to amyloid plaque, tau aggregation, cell death, and synapse loss, and to investigate the spatial relationships between the above localized pathologies and cellular responses. Successful development and implementation of this technology would be highly informative for AD research.

Spatially resolved transcriptomic technologies are capable of mapping single-cell transcriptomic profiles within tissue architecture^28–32^, but many existing ones are incompatible with protein detection in the same tissue sections or limited by the spatial resolution and/or gene coverage. For example, a recent study uncovered amyloid plaque-induced genes (PIG) using Spatial Transcriptomics with fluorescent staining of adjacent brain sections to correlate the positions of plaques with local gene expression^33^. However, the resolution was limited to 100 µm and only a small set of genes were verified at cellular resolution.

We previously developed an image-based *in situ* RNA sequencing method called STARmap (<u>s</u>patially-resolved transcript <u>a</u>mplicon readout <u>map</u>ping) for single-cell transcriptional state profiling in three-dimensional (3D) intact brain tissues^34, 35^. Here we introduce STARmap with <u>p</u>rotein localization and <u>u</u>nlimited <u>s</u>equencing (STARmap PLUS), enabling simultaneous high-resolution spatial transcriptomics concomitantly with specific protein localization in the same tissue section. We employed STARmap PLUS to draw a comprehensive transcriptomic atlas of AD at 200 nm resolution across all brain cell types during the development of amyloid plaque and tau pathology (Fig. 1a). In this study, we applied STARmap PLUS to TauPS2APP triple transgenic mice, an established mouse AD model that exhibits both amyloid plaque and tau pathologies. TauPS2APP mice express mutant forms of hPresenilin 2 (PS2), hAPP and hTau and show age-related brain amyloid deposition, tauopathy, gliosis, neurodegeneration and cognitive deficits^34, 35^. By mapping a targeted list of 2,766 genes, we created a spatial cell atlas of 8- and 13-month old TauPS2APP mice in the context of extracellular Aβ plaques and intracellular hyperphosphorylated tau accumulation at subcellular resolution. Single-cell transcriptomic analysis identified disease-associated gene pathways across diverse cell types in the cortical and hippocampal regions of the TauPS2APP model in comparison with control samples. Integrating the spatial maps of diverse cell types and states at different disease stages, we propose a comprehensive spatiotemporal model of AD disease progression. These studies provide important tools and resources for mechanistic understanding of AD and other neurodegenerative diseases at cellular and molecular levels.

**Fig. 1.**
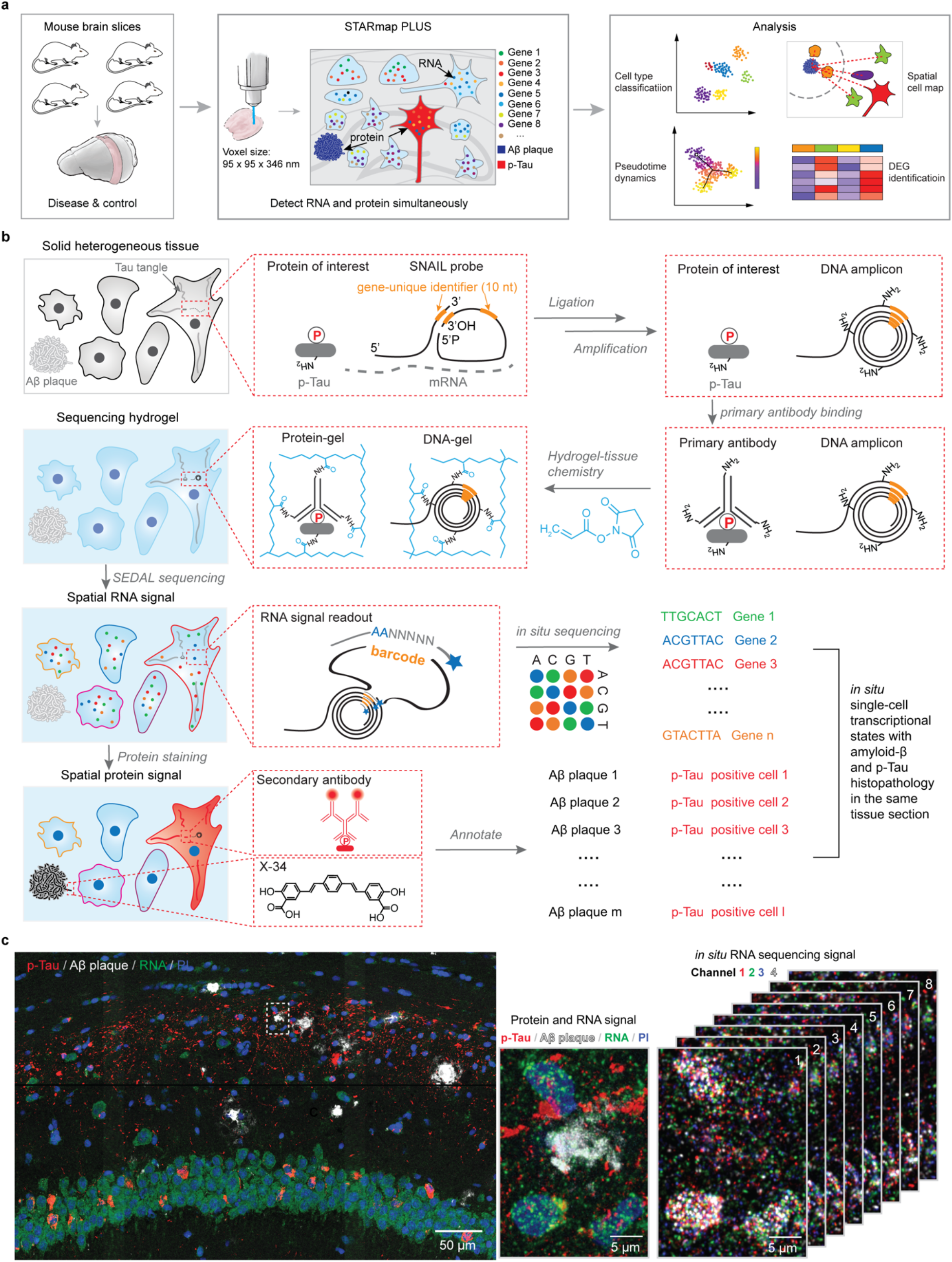
Simultaneous mapping of cell types, single-cell transcriptional states, and tissue histopathology at 200-nm resolution by STARmap PLUS. **a,** Overview of STARmap PLUS, an integrative *in situ* method, capable of simultaneously mapping thousands of RNA species and protein disease markers in the same intact three-dimensional (3D) tissue at subcellular (200 nm) resolution. **b,** Schematics of STARmap PLUS. After the brain tissue is retrieved and fixed, the intracellular mRNAs are targeted by a pair of SNAIL (<u>s</u>pecific <u>a</u>mplification of <u>n</u>ucleic <u>a</u>cids via intramolecular ligation), which are then enzymatically ligated and amplified to generate amine-modified cDNA amplicons *in situ*. Meanwhile, protein markers are labeled with primary antibodies. Next, tissues with amine-modified cDNA amplicons, proteins and primary antibodies are modified by acrylic acid N-hydroxysuccinimide ester (AA-NHS) and copolymerized with acrylamide to generate a hydrogel-tissue hybrid that fixes the locations of biomolecules for *in situ* mapping. Each cDNA amplicon contains a gene-specific identifier sequence (orange) that can be read out through in situ sequencing with error reduction by dynamic annealing and ligation (SEDAL). Lastly, fluorescent protein stainings (secondary antibody and small-molecule dye X-34 stainings) were applied to visualize protein signals. **c,** Representative images showing the simultaneous mapping of cell nuclei, cDNA amplicons, and protein signals in the brain slice from a 13-month old TauPS2APP mouse. Left: The 3D projection of the raw confocal fluorescence image of the CA1 region of the hippocampus. Middle: A zoom-in view of the dashed region in the left panel, which shows the last cycle of tissue histopathology imaging that detects both protein and cDNA amplicon: red, immunofluorescent staining of p-Tau (AT8 primary antibody followed by fluorescent goat anti-mouse secondary antibody); white, X-34 staining of Amyloid β plaque; green, fluorescent DNA probe staining of all cDNA amplicons; and blue, Propidium Iodide (PI) staining of cell nuclei. Right: 8 cycles of *in situ* RNA sequencing of the view in the middle panel; each color represents a fluorescent channel in one round of *in situ* sequencing.

## Results

### Method development of STARmap PLUS

The original STARmap method is incompatible with histological staining (immuno-or chemical staining) and limited to detecting 1,024 genes^32^. To overcome such limitations, in STARmap PLUS, we developed the experimental protocol to incorporate antibody staining (in this study, AT8 antibody, detecting hyperphosphorylated tau) and dye staining (X-34, detecting Aβ plaque) into the library preparation and *in situ* sequencing steps (Fig. 1b, Extended Data Fig. 1a-b). In brief, STARmap PLUS entails the following steps: 1) mRNAs within fixed brain sections are detected by a pair of SNAIL (<u>s</u>pecific amplification of <u>n</u>ucleic <u>a</u>cids via intramolecular ligation) probes (Fig.1b), and enzymatically amplified as cDNA amplicons; 2) specific proteins of interest are labeled with primary antibody; 3) the cDNA amplicons, primary antibodies, and endogenous proteins (*e.g.*, plaques and tau) are chemically modified and copolymerized in a hydrogel matrix; 4) each cDNA amplicon contains a gene-unique identifier (barcode) that is read-out through *in situ* sequencing with error-reduction by dynamic annealing and ligation (SEDAL); 5) fluorescent secondary antibody and small-molecule dye X-34 are then applied to visualize specific proteins and their localization. Besides protein labeling capability, we expanded the gene-coding barcode in the DNA probes from 5 to 10 nucleotides (resulting in 10^6^ coding capacity for STARmap PLUS) that is sufficient to encode more than 20,000 genes (Fig. 1b).

We applied STARmap PLUS to investigate how AD-related pathology, including amyloid deposition and hyperphosphorylated tau, influences brain cell states at the transcriptomic level at single-cell resolution in intact brain tissue in which spatial relationships between protein pathology, cell location and mRNA changes are maintained and readily measured. A curated list of 2,766 genes were extracted from previous bulk and single-cell RNA-seq studies and diverse AD-related databases (Supplementary Table 1)^8, 9, 12, 13, 36–41^. We performed 8 rounds of *in situ* sequencing to map RNAs and 1 round of post-sequencing imaging to locate Aβ plaques and hyperphosphorylated tau (p-Tau) in coronal sections of the brains from TauPS2APP mice and control mice (Fig. 1c and Extended Data Fig. 1c).

Transgenic TauPS2APP mice develop progressive amyloid plaques and tau pathology starting at 4.5 months of age, growing exponentially from 6 to 8 months of age, and rising steadily from 9 months of age. Prominent neurodegeneration, measured by disintegrative staining or volumetric MRI, becomes apparent by 9 months^34, 35^. Thus, brain sections were collected and analyzed at 8 months (when tau and Aβ pathology have set and are expanding) and 13 months (a more advanced disease stage with severe pathology and elevated neuroinflammatory activity)^35^. We focused on the cortex and hippocampus, two of the most susceptible regions in AD. STARmap PLUS revealed that Aβ plaques were prominent in both cortex and hippocampus whereas AT8 p-Tau immunoreactivity was strongest in the CA1 region of the hippocampus (Fig. 1c, Extended Data Fig. 1c), which is consistent with previous reports^34, 35^. A total of 19,932 cells from two TauPS2APP mice and 17,135 cells from two control mice (non-transgenic littermates) were imaged at 200 nm resolution (Fig. 1a). After projecting 3D RNA reads to two-dimensional planes, segmenting cells, and filtering single-cell transcriptional profiles by quality control (see Methods), the remaining 33,106 cells pooled from all 4 samples were subjected to downstream analysis (Extended Data Fig. 1d).

### Hierarchical cell classification and spatial analysis

To identify cell types from the STARmap PLUS data, we adopted a hierarchical clustering strategy, where top-level clustering served to classify cells into common cell types shared by all samples, and sub-level clustering served to further identify disease-associated subtypes. During top-level clustering, the Leiden algorithm was applied to the Uniform Manifold Approximation and Projection (UMAP) representation of all transcriptomic profiles^42, 43^. Through Leiden clustering, we identified 13 major clusters and annotated the cell types according to their spatial distribution and previously reported gene markers^38, 39, 41^(Fig. 2a and Extended Data Fig. 2a). For example, excitatory neurons were annotated by their high expression levels of genes related to ion channels and synaptic signaling such as *Vsnl1*, *Snap25*, and *Dnm1*. Inhibitory neurons were separated by their enrichment of gamma-aminobutyric acid (GABA) transporters *Slc6a1*. Other non-neuronal cell type specific markers such as *Aldoc* for astrocyte, *Bsg* for endothelial cell, *Ctss* for microglia, *Plp1* for oligodendrocyte were used to annotate corresponding clusters (Extended Data Fig. 2a). The UMAP plots of TauPS2APP samples showed differential distribution of cells within the astrocyte, microglia, oligodendrocyte and dentate gyrus (DG) clusters in comparison with controls, suggesting possible disease-associated cell subtypes (Fig. 2a). Thus, we further investigated the transcriptomic heterogeneity within each major cell type, which further identified 27 sub-level clusters based on their transcriptomic signatures (Fig. 2b).

**Fig. 2.**
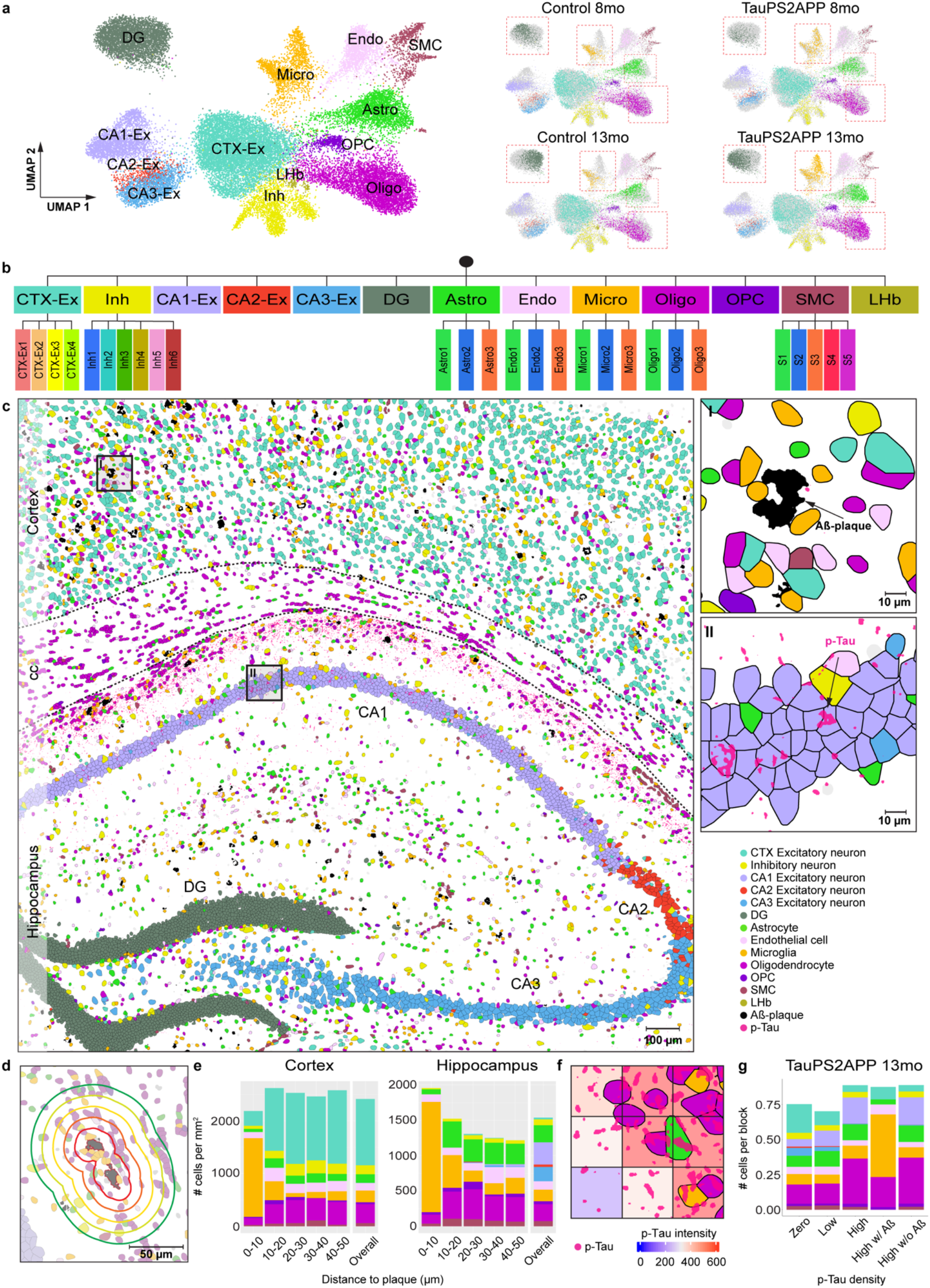
Top-level cell-type classification and spatial analysis in the brain slices of TauPS2AAPP and control mice. **a,** Uniform Manifold Approximation and Projection (UMAP) plot visualization of transcriptional profiles of 33,106 cells collected from coronal brain sections of TauPS2APP and control mice at 8 and 13 months. 13 major cell types were identified using the Leiden clustering. Red dashed boxes highlight the clusters of cells that show a significant difference between TauPS2APP and control samples in the UMAPs. **b,** Hierarchical taxonomy of cell types showing the 13 top-level clusters: Cortex excitatory neuron (CTX-Ex, 8,687 cells), Inhibitory neuron (Inh, 2,005 cells), CA1 excitatory neuron (CA1-Ex, 2,754 cells), CA2 excitatory neuron (CA2-Ex, 436 cells), CA3 excitatory neuron (CA3-Ex, 1,878 cells), Dentate Gyrus (DG, 4,377 cells), Astrocyte (Astro, 2,884 cells), Endothelial cell (Endo, 1,849 cells), Microglia (Micro, 1,723 cells), Oligodendrocyte (Oligo, 4,966 cells), Oligodendrocyte precursor cell (OPC, 549 cells), Smooth muscle cell (SMC, 877 cells), Lateral Habenula neuron (LHb, 121 cells) and 27 sub-level clusters (subclusters) identified during the cell type classification process according to their representative gene markers. The transcriptional profile of each interested top-level cluster was analyzed using Leiden clustering again to identify subclusters. **c,** Representative spatial cell-type atlas with Aβ and tau pathologies in cortical and hippocampal regions of TauPS2APP 13-month sample. Top-level cell types were color-coded based on the legend (inset). Aβ plaque and p-Tau protein were colored as black and magenta, respectively. The imaging area was separated into Cortex, Corpus Callosum (*CC*), and Hippocampus manually with prior knowledge and the boundaries were marked by a black dash lines. Scale bar, 100 µm. Zoom-in sections: (I) zoom-in section in the cortical region showing an Aβ plaque surrounded by different types of cells; (II) zoom-in section in the hippocampal region with p-Tau protein signal; Scale bars, 10 µm. **d,** Schematics illustrating the spatial patterns analysis of cell type compositions around the Aβ plaque. 5 concentric boundaries that are 10, 20, 30, 40, 50 µm from each plaque were generated to quantify the cell-type composition of each layer. When overlapping, two stripes were merged to prevent repetitive counting. Scale bar, 50 µm. **e,** Representative spatial distribution of cell type compositions around Aβ plaque for TauPS2APP 13-month sample. Stacked bar plot showing the density (cell count per mm^2^) of each major cell type at different distance intervals (0-10, 10-20, 20-30, 30-40, 40-50 µm) to the Aβ plaque. The cell density of each major cell type in each area was included as the reference for comparison. **f,** Schematics illustrating the method used for p-Tau signal quantification. Tissue sections were divided into a grid of 20 µm x 20 µm. The color represented integrated intensity of p-Tau in each square and was used as a metric to analyze the extent of colocalization of p-Tau with different cell types. **g,** Cell-type composition analysis based on the 20 µm x 20 µm grid in the TauPS2APP sample at 13 months ranked by p-Tau density. The blocks divided by the grid lines were ranked by the percentage of Tau positive pixels and grouped into 3 bins: 0% (zero p-Tau), 1-50% (low p-Tau), 51-100% (high p-Tau). The high p-Tau group was further divided by plaque positive versus negative groups to dissect the influence of plaque and tauopathy on cell-type distribution. Stacked bar plot showing the average number of cells per block for each major cell type.

Because spatial information of RNAs was preserved at subcellular resolution, we were able to generate a high-resolution spatial cell atlas of the cortex and hippocampus in conjunction with precise localization of AD-related histopathology (Fig. 2c and Extended Data Fig. 2b,c). All spatial cell atlases of TauPS2APP and control non-transgenic mice showed similar anatomic structure with major cell-type distribution in cortical and hippocampal regions, confirming the robustness, reproducibility and reliability of top-level clustering results. In TauPS2APP mice, the percentage of plaque area in the whole tissue (cortex and hippocampus) rose from 0.17% (73 Aβ plaques identified with an average size of 89 ± 44.9 µm^2^) in 8-month mouse brains to 0.54% (151 plaques identified with an average size of 157 ± 110.7 µm^2^) in 13-month mouse brains. Meanwhile, no plaques were visible in non-transgenic controls (Extended Data Fig. 1c). p-Tau signal was observed mostly in the hippocampal region of the TauPS2APP mice at 8 months and became more pronounced at 13 months in both hippocampal and cortical regions (Fig. 2c and Extended Data Fig. 2c).

How are cell distributions and cell states affected by the nearby presence of amyloid plaque and tauopathy? First, given the different cell-type compositions in different parts of the brain, we analyzed the cell distributions and states in the cortex and hippocampus separately. Next, to quantify cell-type distribution in the spatial relationship to Aβ plaque, we calculated the density of different cell types at the first five 10 µm intervals surrounding plaques (Fig. 2d, Methods). Among the 13 major cell types, microglia, astrocytes, oligodendrocytes, and oligodendrocyte precursor cells (OPC) showed relative enrichment in the vicinity of plaques in comparison with the overall density of that cell type in the cortex and hippocampus in TauPS2APP mice (Fig. 2e and Extended Data Fig. 2d). The most striking change was detected in microglia: they were the most prevalent cell type in the 10 µm ring around the plaque, where they were markedly concentrated compared with the normal density in that brain region and often seemed to be in direct contact with the plaques. Astrocytes, oligodendrocytes and OPC showed modest but significant enrichment at the 10-30 µm distance compared to their overall average density but were depleted in the immediate neighborhood of plaques (<10 µm), perhaps crowded out by microglia. Both excitatory and inhibitory neurons were also depleted within 10 µm around Aβ plaques in comparison with the overall density of each neuron type.

In contrast to the localized macroaggregates of Aβ plaques, p-Tau (AT8) immunoreactivity was dispersed throughout neuronal cell bodies and axon bundles. To investigate the spatial correlation of p-Tau intensity with different cell types, we divided the spatial atlas into blocks of 20 µm-spaced grids (Fig. 2f) and analyzed the covariation of cell-type composition versus p-Tau density. Oligodendrocytes were found enriched in the blocks with high p-Tau intensity regardless of the presence or absence of Aβ plaques (Fig. 2g).

The top-level cell clustering and spatial analyses revealed that microglia, astrocytes, OPCs, oligodendrocytes, and neuronal cells showed the biggest changes in their transcriptional profiles, spatial distributions, or both. These cell types were thus selected for in-depth sub-clustering analyses to pinpoint disease-associated subtypes, cell states, and gene pathways in the following sections (Figs. 2b and 3-6).

### Disease-associated microglia directly contact Aβ plaques from early disease stage

We first investigated the heterogeneity within the microglia population by sub-clustering analysis of transcriptomic profiles. Three subpopulations were identified and categorized as Micro1 (45%), Micro2 (24%), and Micro3 (31%) (Fig. 3a). Micro1 and Micro2 subtypes were present in both TauPS2APP and control samples, while the Micro3 subpopulation cells were almost absent in controls and expanded greatly from 8 months to 13 months in the TauPS2APP brains (Fig. 3a). Micro1 and Micro2 both express the marker *P2ry12* and may correspond to subtypes of homeostatic microglia^10^: Micro1 expresses high levels of *P2ry12* whereas Micro2 shows downregulation of *P2ry12* and upregulation of *Tgfbr1* and *Gpr34*. The Micro3 subtype expressed high levels of *Cst7*, *Ctsb*, *Trem2*, and *Apoe*, which are characteristic of DAM and associated with neurodegeneration^10^ (Fig. 3b). Given their strong association with plaques in the TauPS2APP disease model and expression of known DAM gene markers, we believe that Micro3 microglia defined in STARmap PLUS are equivalent to DAM microglia previously described in conventional single cell RNA-seq studies^10, 16^.

**Fig. 3.**
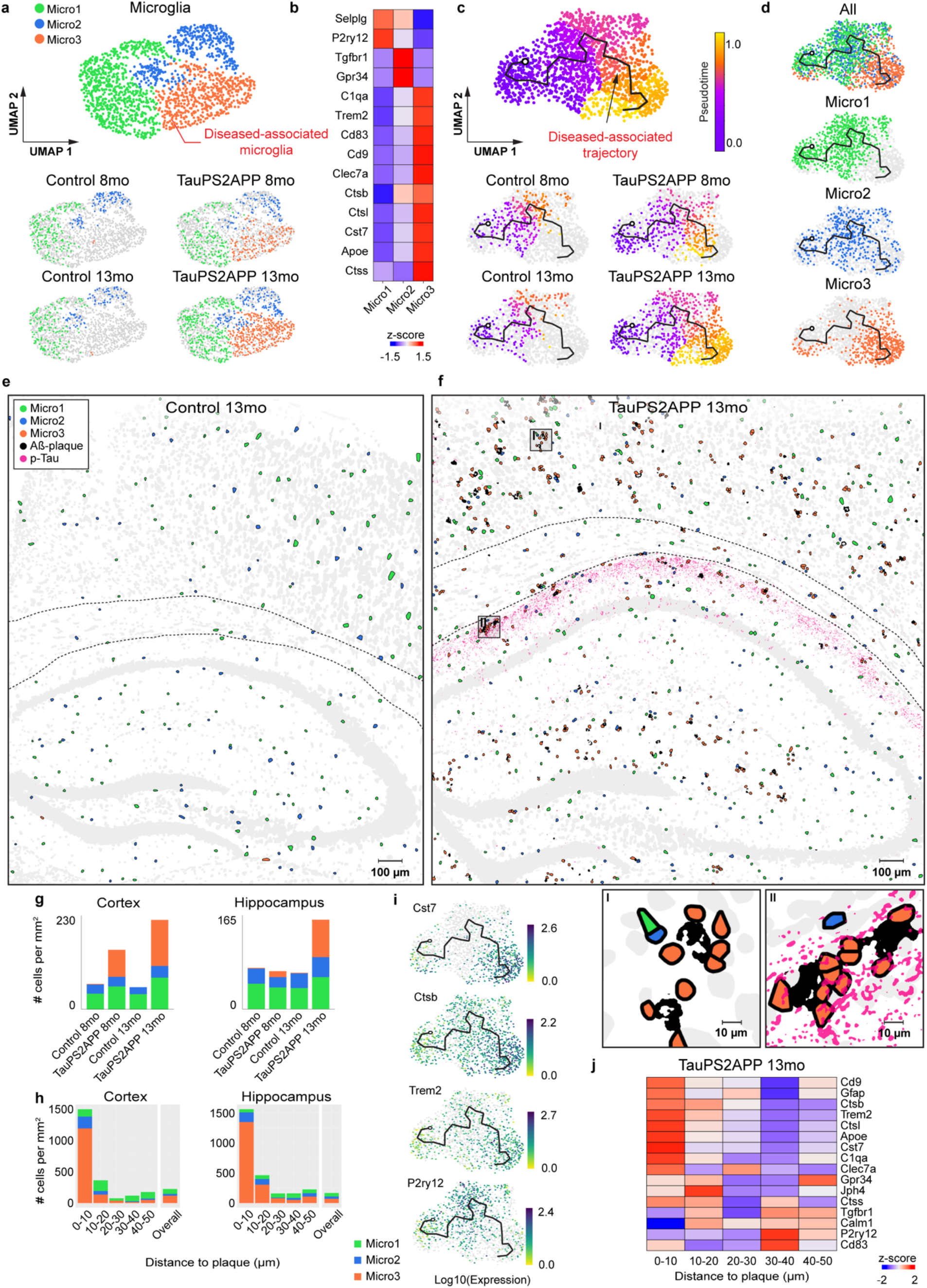
Spatiotemporal gene expression analysis of microglia in TauPS2APP and control samples. **a,** Subclustering of microglia cell population. Top: Zoom-in UMAP visualization of 1,723 microglia cells from Fig. 2a showing three subclusters of microglia identified by the Leiden clustering: Micro1 (n = 779), Micro2 (n = 415), and Micro3 (n = 529). The Micro3 cluster was considered as a disease-associated microglia (DAM) population by its gene markers according to previous reports and significant enrichment in diseased samples^10^. Bottom: UMAP visualization of subclusters of microglia populations for each sample (Control 8 month, TauPS2APP 8 month, Control 13 month, TauPS2APP 13 month). **b,** Expression level of representative gene markers among different subclusters colored by row-wise z-score. **c,** UMAP pseudotime trajectory visualization of microglia population. Top: UMAP visualization of pseudotime trajectory of microglia population generated by Monocle 3. Colormap represents the pseudotime value. A corresponding pseudotime trajectory was plotted on the UMAP embedding. Trajectory starting anchor was manually selected based on the control sample at 8 months. Bottom: Cells from different samples were highlighted separately in UMAP embedding. **d,** UMAP plots showing different types of microglia identified in (**a**) in pseudotime embedding. **e,f,** Spatial cell maps of microglia in the 13-month control and TauPS2APP samples. Scale bar, 100 µm. Insets show zoom-in regions highlighted by black boxes (I, II). Scale bar, 10 µm. Dashed black lines mark the boundaries between cortex, corpus callosum and hippocampus. **g,** Stacked bar chart showing the density (count per mm^2^) of each microglia sub-cluster in the cortex and hippocampus region for each sample. The colors in the bar plots correspond to the cell type legend in (**e**). The plots show a significant enrichment of the DAM in both TauPS2APP samples at 8 and 13 months. **h,** Stacked bar chart showing the density (count per mm^2^) of each microglia sub-cluster in each distance interval (0-10, 10-20, 20-30, 30-40, 40-50 µm) centered by the Aβ plaque at 13 months. The overall cell density of each subpopulation in each area was included as the reference for comparison (Overall). **i,** DEGs on pseudotime embedding. UMAPs showing the expression of four representative gene markers of microglia subtypes on the pseudotime embedding. The color indicates the log10(mean gene expression value) of the gene in each cell. **j,** Matrix plot showing the z-scores of top DEGs and representative markers of microglia across multiple distance intervals (0-10, 10-20, 20-30, 30-40, 40-50 µm) from plaques.

It is noteworthy that clustering analysis effectively identifies distinct cell subtypes but does not capture well multi-step cell-state transitions. To uncover cell-state transitions during disease progression and determine the relationship among different subtypes, we deployed Monocle pseudotime analysis^44^, a widely used computational tool for reconstructing cell differentiation trajectory, as a complement to subtype analysis in the following sections. We aimed to reconstruct the presumptive path along which microglia alter their state by pseudotime trajectory analysis of the single cell transcriptomic profiles (Fig. 3c). By this computational approach, the microglia population showed a linear pseudotime trajectory that aligned well with the real disease progression timeline: microglia in control mice were enriched at the starting point of the trajectory while those in TauPS2APP mice shifted along the trajectory from 8 months to 13 months (Fig. 3c). Along the trajectory, Micro1 and Micro3 cells are distinctively distributed at the early and late pseudotime values, respectively, whereas Micro2 cells are more intermediate in pseudotime position and blended into Micro1 and Micro3 populations (Fig. 3d).

Do the three microglia subtypes Micro1-3 show different spatial patterns? In the cortex, all three subtypes showed increased cell density at both ages of TauPS2APP mice, while in the hippocampus, the density of Micro2 and Micro3 increased at 13 months (Fig. 3e-g). With the cell-type composition analysis around Aβ plaques, we found: (i) Micro3 is the most predominant cell type (>70%) within the 10 µm ring around plaque (Fig. 3h and Extended Data Fig. 3b); (ii) Micro2 cells were enriched within 10 µm distance around the plaques in both the cortex and hippocampus, while Micro1 showed enrichment close to plaque only in the cortex (Fig. 3h, Extended Data Fig. 3b and Supplementary Table 2); (iii) from 8 months to 13 months, the density of Micro3 drastically increased within the 10 µm ring near plaque, whereas the local cell density of Micro1 and Micro2 near plaque (<10 µm) remained the same or decreased (Fig. 3h and Extended Data Fig. 3b, Supplementary Table 2); (iv) the pseudotime distribution of microglia around plaque showed that the pseudotime values of microglia cells within 10 µm were higher than those of cells far away from plaques and these pseudotime values increased from 8 months to 13 months (Extended Data Fig. 3c,d). Together, the spatial analysis of microglia cell subtypes and pseudotime distribution suggest that the extent of microglia activation strongly correlates with their close association with plaques (<10 µm), suggesting a local transition from Micro1 and Micro2 to Micro3 near plaque along disease progression.

To get a more comprehensive understanding of microglia response to AD pathology at the molecular level, we analyzed differential expressed genes (DEGs) of microglia along disease progression and in relation to plaques (Fig. 3i,j and Extended Data Fig. 3e,f, Supplementary Table 3). Most of the up-regulated microglial DEGs identified in the TauPS2APP mouse brains overlap with Micro3 (DAM) gene markers and with up-regulated “spatial DEGs” near plaques; these genes are involved in biological processes such as regulation of immune response (i.e. *Apoe*, *C1qa*, *Cd74*), regulation of cell activation (i.e, *Cd9*, *Cd83*, *Cst7*), cellular protein catabolic process (i.e. *Ctss*, *Ctsb, Sumo2, Cul1*) (Extended Data Fig. 3e,f). Notably, relating to a recent study of human AD patient samples^45^, our study revealed that a few upregulated genes (i.e. *Ccl3, Trem2, Csf1r*) in the DAM microglia of the TauPS2APP model may be mediated by the activation of ERK1/ERK2 cascade (Extended Data Fig. 3f).

### Disease-associated astrocytes emerge near the plaque-DAM complex at a later stage

Astrocyte was another non-neuronal cell type that showed a significant difference in TauPS2APP versus control. Sub-clustering analysis of the astrocytes identified three transcriptionally distinct subpopulations Astro1 (37%), Astro2 (44%), and Astro3 (19%) (Fig. 4a). Astro3 cells were rarely found in control mice but the Astro3 subpopulation greatly expanded from 8 months to 13 months in TauPS2APP mice (Fig. 4a). Transcriptomic changes in Astro3 astrocytes closely resembled those of disease-associated astrocytes (DAA) previously described in AD models, characterized by upregulation of genes such as *Gfap*, *Vim*, *Apoe* (Fig. 4b)^9^. Based on this transcriptomic similarity and their association with TauPS2APP mice we annotated the Astro3 subtype as DAA. Astro1 and Astro2 are present in both control and TauPS2APP mice; their gene markers correspond with previously reported low- and intermediate-*Gfap* cell populations^9^. In comparison with controls, the overall cell density of Astro1 declined in the TauPS2APP mice whereas the density of Astro2 increased in the cortex.

**Fig. 4.**
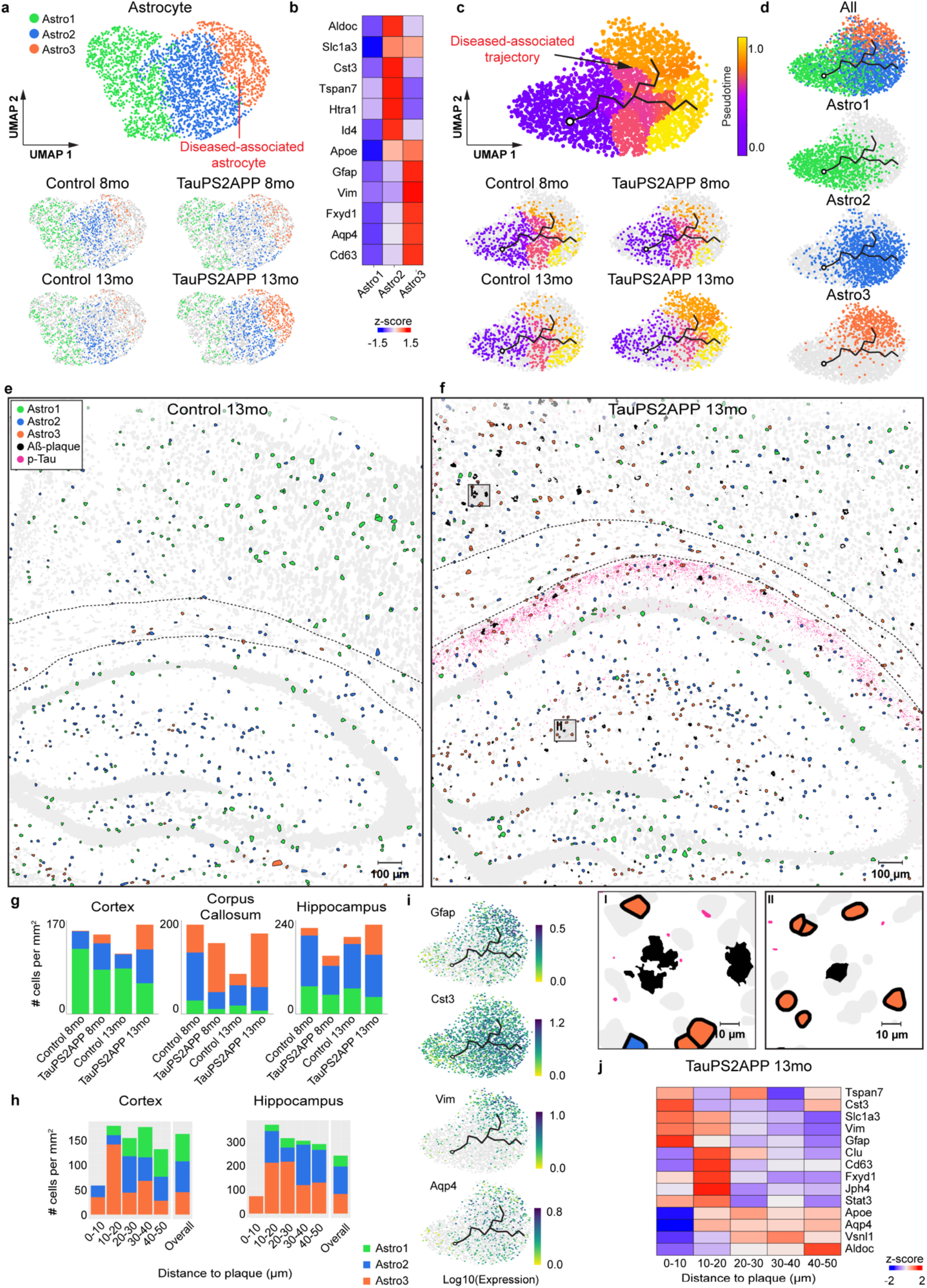
Spatiotemporal gene expression analysis of astrocytes in TauPS2APP and control samples. **a,** Subcluster of astrocyte cell population. Top: Zoom-in UMAP visualization of 2,884 astrocytes identified from Fig. 2a showing three subclusters of astrocytes identified by the Leiden clustering: Astro1 (n = 1,068), Astro2 (n = 1,271), and Astro3 (n = 545). The Astro3 cluster was annotated as disease-associated astrocyte (DAA) by its gene markers according to previous reports^9^ and enrichment in diseased samples. Bottom: UMAP plots of subclusters of astrocytes for each sample (Control 8 month, TauPS2APP 8 month, Control 13 month, TauPS2APP 13 month). **b,** Expression level of representative markers across different sub-clusters. Colored by row-wise z-score. **c,** UMAP pseudotime trajectory visualization of astrocyte cell population. Top: UMAP visualization of pseudotime trajectory of astrocytes generated by Monocle 3. Colormap represents the pseudotime value. A corresponding pseudotime trajectory was overlapped on the UMAP embedding. Trajectory starting anchor was manually selected based on the Astro1 population. Red arrow highlights bifurcation point on the trajectory related to the disease-associated gene expression changes. Bottom: Cells from different samples were highlighted separately in UMAP embedding. **d,** UMAP plots showing different types of astrocytes identified in (**a**) in pseudotime embeddings. **e,f,** Spatial cell map of astrocyte population in 13-month control and TauPS2APP samples. Scale bar, 100 µm. Insets show the zoom-in regions highlighted by black boxes (I, II). Scale bar, 10 µm. Dashed black lines mark the boundaries between cortex, corpus callosum and hippocampus. **g,** Stacked bar chart showing the density (count per mm^2^) of each astrocyte sub-cluster in the cortex, Corpus Callosum and Hippocampus region for each sample. The colors in the bar plots correspond to the cell type legend in (**e**). The plots show a significant enrichment of the DAA in the cortex, especially at 13 months. **h,** Stacked bar chart showing the density (count per mm^2^) of astrocyte sub-cluster in each distance interval (0-10, 10-20, 20-30, 30-40, 40-50 µm) around the Aβ plaque at 13 months. The cell density of each subpopulation in each area was included as the reference for comparison (Overall). **i,** DEGs on pseudotime embedding. UMAPs showing the expression of four disease up-regulated genes on the pseudotime embedding, the color scale of the raw counts was adjusted by log10 transformation. **j,** Matrix plot showing the z-scores of top DEGs and representative markers of astrocytes across multiple distance intervals (0-10, 10-20, 20-30, 30-40, 40-50 µm) from plaques.

Despite the apparent linear gradient of many ‘marker’ genes across Astro1-3 subtypes, we noted a bifurcation path based on pseudotime trajectory analysis of the astrocyte population (Fig. 4c). By visualizing the subtype annotation along the pseudotime trajectory, we identified that the longest path (lower path) matches with the gradient from Astro1 (the starting point) and Astro 2 (the ending point) shared by TauPS2APP and control samples. In contrast, the short branch from the bifurcation point (labeled as disease-associated trajectory in Fig. 4d) is enriched with Astro3 (DAA) from the 13-month TauPS2APP sample and represented the transition from non-diseased states (mostly Astro2) to Astro3.

The spatial cell map of astrocyte subtypes showed that Astro1 cells are preferentially located near the cell bodies of cortical and hippocampal neurons, whereas Astro2 cells are more concentrated in corpus callosum, hippocampal neuropil layer, and stratum lacunosum-moleculare (Fig. 4e-g and Extended Data Fig. 4a). Cell-type analysis in relation to tissue pathology revealed that while DAMs closely surround plaque (<10 µm), Astro3 (DAA) cells were enriched around plaque at an intermediate distance (10-20 µm in the cortex, 10-30 µm in the hippocampus) in TauPS2APP mice at 13 months (2-3 fold increase in Astro3 density in this intermediate shell versus >40 µm from plaques) (Fig. 4h, Supplementary Table 2). Astro2 was also enriched near the plaques (10-20 µm) as the major astrocyte subtype at 8 months, however, they were outnumbered by Astro3 at 13 months (Fig. 4h, Extended Data Fig. 4b and Supplementary Table 2). The observed shift of the astrocyte population around plaque from Astro2 to Astro3 from 8 to 13 months, in combination with the disease-associated pseudotime trajectory from Astro2 to Astro3 (Fig. 4d), suggests that there might be a conversion of Astro2 to Astro3 (DAA) near plaques during disease progression. Spatial analysis of the astrocyte pseudotime distributions near plaques also shows a shift from a mixture of low-middle values to a more homogeneous astrocyte population with high pseudotime values from 8 to 13 months, especially in the 10-20 µm shell (Extended Data Fig. 4c,d).

Most of the DEGs from the astrocytes of TauPS2APP versus control samples were related to glial cell differentiation and gliogenesis (or astrogliosis, i.e. *Gfap*, *Vim*, *Clu*, *Stat3*) (Extended Data Fig. 4e,f and Supplementary Table 3). The gene expression profiles of top DEGs identified on the pseudotime embedding showed that the expression pattern of *Vim* best correlates with the DAA population, whereas the molecular gradients of *Gfap* resembled the disease-associated trajectory from Astro 2 to Astro 3 (Fig. 4i). Spatial analysis of gene expression around plaques in the 13-month TauPS2APP mice further support the spatial enrichment of DAA near plaques (Fig. 4j).

### OPCs are enriched in the intermediate vicinity of Aβ plaques and oligodendrocyte subtypes infiltrate the hippocampal alveus colocalizing with p-Tau

Sub-clustering analysis identified four subtypes within the oligodendrocyte lineage (Fig. 5a,b): Oligo1 (78%), Oligo2 (3%), Oligo3 (9%), and OPC (10%). The oligodendrocyte gene marker *Plp1* marks all three oligodendrocyte subtypes (Oligo1-3) whereas differential expression of other genes such as *Klk6* and *Cldn11* mark Oligo2 and Oligo3 populations, respectively (Fig. 5b). All four subtypes are present in TauPS2APP and control brains, though the abundance of Oligo2 and Oligo3 oligodendrocytes increased by a factor of 2-3 (Supplementary Table 2) in response to amyloid and tau pathology at 13 months.

**Fig. 5.**
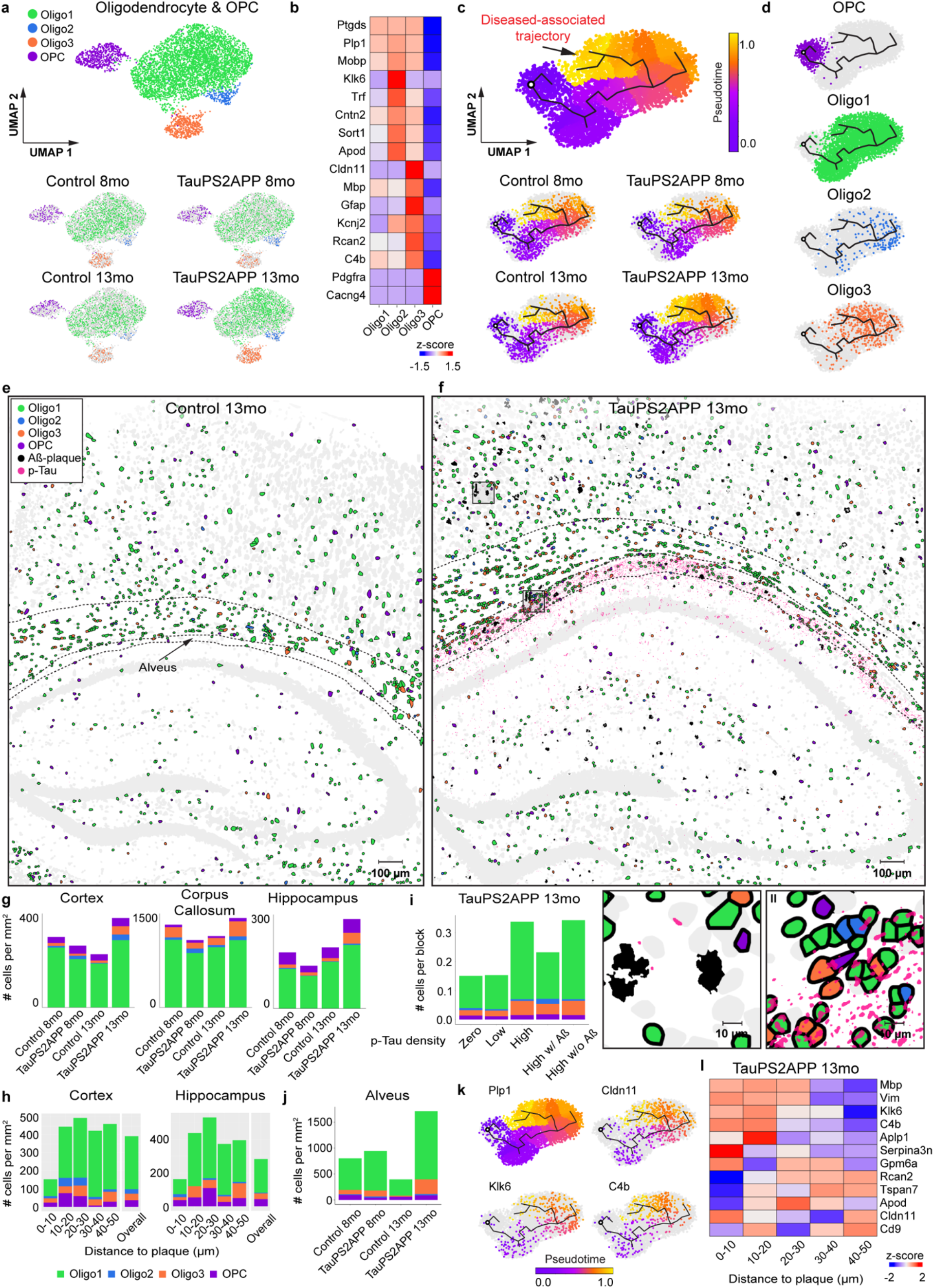
Spatiotemporal gene expression analysis of oligodendrocytes and precursor cells in TauPS2APP and control samples. **a,** Subcluster of Oligodendrocytes and precursor cell (OPC) population. Top: Zoom-in UMAP visualization of 4,966 oligodendrocytes and 549 OPCs identified from Fig. 2a showing three subclusters of oligodendrocytes identified by the Leiden clustering: Oligo1 (n = 4,295), Oligo2 (n = 181), Oligo3 (n = 490) and OPC (n = 549). Bottom: UMAP plots of subclusters of oligodendrocytes and OPCs for each sample (Control 8 month, TauPS2APP 8 month, Control 13 month, TauPS2APP 13 month). **b,** Expression level of representative markers across different sub-clusters of oligodendrocytes colored by row-wise z-score. **c,** UMAP pseudotime trajectory visualization of oligodendrocyte related cell population. Top: UMAP visualization of pseudotime trajectory of oligodendrocytes and OPCs generated by Monocle 3. Colormap represents the pseudotime value. A corresponding pseudotime trajectory was plotted on the UMAP embedding. Trajectory starting anchor was manually selected based on the OPC population. Red arrow highlights bifurcation point on the trajectory related to the disease-associated gene expression changes. Bottom: Cells from different samples were highlighted separately in UMAP embedding. **d,** UMAP plots showing different types of oligodendrocytes and OPCs identified in (**a**) in pseudotime embeddings. **e,f,** Spatial cell map of oligodendrocyte related population in 13-month control TauPS2APP samples. Scale bar, 100 µm. Insets show the zoom-in regions highlighted by black boxes (I,II). Scale bar, 10 µm. Dashed black lines mark the boundaries between cortex, corpus callosum, hippocampus and alveus. **g,** Stacked bar chart showing the density (count per mm^2^) of each oligodendrocyte sub-types and OPC in both the cortex, corpus callosum and hippocampus region for each sample. The colors in the bar plots correspond to the cell type legend in (**e**). **h,** Stacked bar chart showing the density (count per mm^2^) of each oligodendrocyte sub-cluster and OPC in each distance interval (0-10, 10-20, 20-30, 30-40, 40-50 µm) around the Aβ plaque at 13 month. The overall cell density of each subpopulation in each area was included as the reference for comparison (all). **i,** Cell-type composition analysis of oligodendrocyte lineages in relation to p-Tau pathology. The tissue region was divided by the 20 µm x 20 µm grid in the TauPS2APP sample at 13 months ranked by p-Tau density. The blocks divided by the grid lines were ranked by the percentage of p-Tau positive pixels and grouped into 3 bins: zero (0%), low (1%-50%), high (50%-100%). The high p-Tau bin is further divided into two groups based on presence or absence of Aβ plaques. Stacked bar plot showing the average number of cells per block for each oligodendrocyte subtype. **j,** Cell density and subtype composition of oligodendrocyte and OPC in the hippocampal alveus region. **k,** Pseudotime embedding of cells expressing marker genes. UMAPs showing the pseudotime distribution of cells expressing representative markers. **l,** Matrix plot showing the z-scores of top DEGs and representative markers of oligodendrocytes across multiple distance intervals (0-10, 10-20, 20-30, 30-40, 40-50µm) from plaques.

The pseudotime trajectory analysis of the combined population of OPC and oligodendrocyte cells recapitulated the known differentiation path from OPC to mature oligodendrocytes (Fig. 5c, lower trajectory). Similar to astrocytes, a disease-associated trajectory diverged from the main path from OPC to oligodendrocytes (Fig. 5c, upper branch). Oligodendrocytes from 13-month TauPS2APP brains were enriched around the disease-associated trajectory branch when compared with 8-month TauPS2APP and controls (Fig. 5c). The disease-associated trajectory contained all three Oligo1-3 subtypes but with an enrichment of Oligo3 cells (Fig. 5d).

Around amyloid plaques in TauPS2APP mice, OPCs were enriched in the 10-30 µm ring at both 8 and 13 months, accumulating to a 40%∼140% higher density than its overall density (Fig. 5e-h and Extended Data Fig. 5a,b and Supplementary Table 2). Oligo1 is the dominant (>70%) oligodendrocyte subtype around plaques among all oligodendrocytes. At 8 months, Oligo1-3 did not show a statistically significant enrichment around amyloid plaques (Extended Data Fig. 5a,b and Supplementary Table 2). At 13 months, the density of Oligo1 increased by 58%-80% at the 10-30 µm distance from the plaques in TauPS2APP mice in comparison with the overall density (Fig. 5h); the average densities of Oligo2 and Oligo3 also increased by 1.5-3 fold at the 10-30 µm distance from plaques but they were only sparsely present near (10-30µm) 13-14% of all plaques. Spatial analysis of the pseudotime values of oligodendrocytes lineages (mostly Oligo1) showed a global upregulation from 8 months to 13 months regardless of their distance to plaques (Extended Data Fig. 5c-e), indicating that the shift of cell states in oligodendrocytes is not spatially confined near plaques.

Given that the density of oligodendrocytes is positively correlated with the density of p-Tau (Fig. 2g), we sought to pinpoint which oligodendrocyte subtype is spatially associated with tauopathy. Using the aforementioned grid-based spatial correlation analysis, we found that in the TauPS2APP mice, the cell density of Oligo1 and Oligo3 increased by 1.5-3 fold in regions with higher p-Tau signals regardless of Aβ pathology (Fig. 5i and Extended Data Fig. 5f). In contrast, the cell density of OPC and Oligo2 increased by 30% and 140% respectively only in the presence of plaques, suggesting that these oligodendrocyte lineage populations respond to amyloid rather than Tau pathology. We observed that p-Tau signals were concentrated in the alveus of the hippocampus where axon bundles of hippocampal neurons course (alveus, Extended Data Fig. 5g). Compared with 13-month control samples, the Oligo1 and Oligo3 densities increased markedly (2-4-fold) in the alveus region of 13-month TauPS2APP mice (Fig. 5j). In total, the spatial analysis of cells of the oligodendrocyte lineage in relation to p-Tau revealed a strong association between tauopathy and the accumulation of Oligo1 and Oligo3 subtypes in the hippocampus alveus.

Through analyzing the DEGs in oligodendrocytes along disease-associated pseudotime trajectory and around plaques, as well as DEGs from the TauPS2APP mice versus control mice, we identified and verified a group of genes (such as *Klk6*, *C4b, Cd9, Serpina3n*) that were strongly upregulated in 13-month old TauPS2APP mice compared to the same age control mice (Supplementary Table 3). Interestingly, oligodendrocytes near plaques exhibited increased expression of these genes, suggesting subpopulation(s) of oligodendrocytes (with elevated expression of *C4b*, *Serpina3n*, *Cd9*) might be responding to plaques or interacting with other cells or cellular compartments (microglia, astrocytes and/or dystrophic neurites) that are affected by amyloid pathology (Fig. 5l). In contrast, the fold change of DEGs between 8-month old TauPS2APP and control mice was less significant (Supplementary Table 3). GO term analysis indicated involvement of oligodendrocyte DEGs in cytokine production (i.e. *Ndrg2, Serpinb1b*) and regulation of synaptic plasticity (i.e. *Jph4, App, Nrgn*) during disease progression.

### Susceptibility of different neuronal types to Aβ plaques and p-Tau

Besides cellular changes in glial cells, the transcriptomic responses in neurons are critical for understanding the mechanisms of neurodegeneration. Sub-clustering analysis of the neurons in cortex and hippocampus identified eight excitatory neuron subtypes and six inhibitory neuron subtypes. As visualized in the spatial cell map of neurons (Fig. 6a,b and Extended Data Fig. 6a-d), the four subtypes of cortical excitatory neurons correspond to different cortical layers (CTX-Ex1 corresponds to layers 2/3, CTX-Ex2/3 corresponds to layers 4/5, CTX-Ex4 corresponds to layer 6.); the excitatory neuronal types in the hippocampal region correspond to the principal cells of DG, CA1, CA2 and CA3. Among the four subtypes of inhibitory neurons, *Pvalb* and *Sst* neurons were present in higher density in cortex while *Cnr1* and *Lamp5* neurons were more abundant in the hippocampus (Extended Data Fig. 6d).

**Fig. 6.**
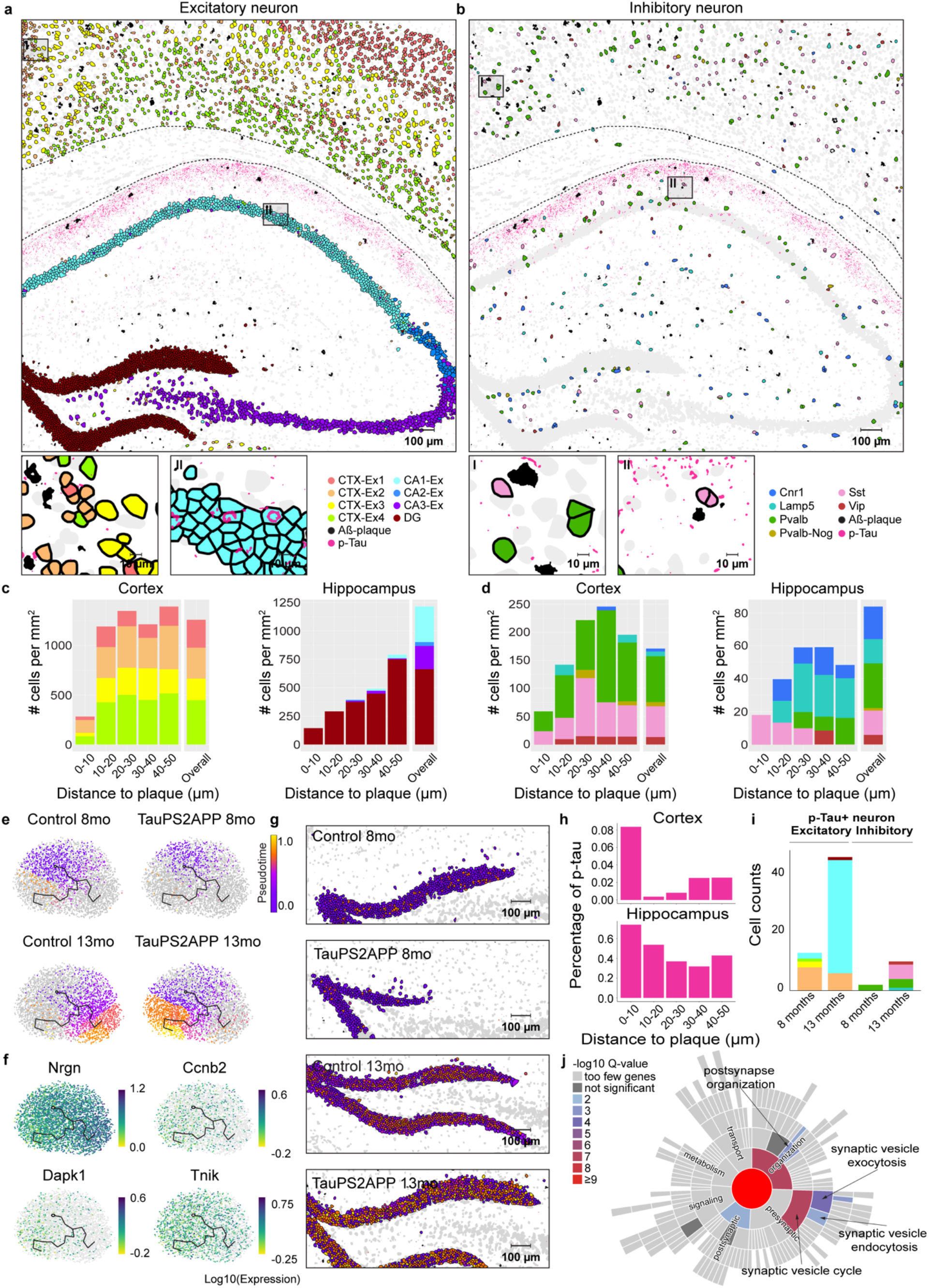
Spatiotemporal gene expression analysis of neurons in TauPS2APP and control samples. **a,b,** Top: Spatial map of Aβ plaque and p-Tau with excitatory (**a**) and inhibitory (**b**) neuron in the TauPS2APP 13 month sample. Scale bar, 100 µm. Total cell counts: CTX-Ex1: 2,292; CTX-Ex2: 2,766; CTX-Ex3: 1,184; CTX-Ex4: 2,445; CA1: 2,754; CA2: 436; CA3: 1,878; DG: 4,377; Cnr1: 224; Lamp5: 196; Pvalb: 831; Pvalb-Nog: 74, Sst: 508. Vip: 172. Bottom: high magnification views of areas indicated in the black boxes on the top panel. Scale bar, 10 µm. **c,d,** Stacked bar charts showing the density (count per mm^2^) of each subcluster of excitatory (**c**) and inhibitory (**d**) neuron population from different brain regions at different distance intervals (0-10, 10-20, 20-30, 30-40, 40-50 µm) to the Aβ plaque of the 13-month TauPS2APP sample. The overall cell density of each subpopulation in each area was included as the reference comparison standard (Overall). **e,** UMAP of dentate gyrus (DG) cell population labeled with pseudotime trajectory showing divergent paths of TauPS2APP and control samples at 13 months. Colormap represents the pseudotime value. Trajectory starting anchor was manually selected based on the control sample at 8 months and labeled by the black circles. **f,** UMAPs showing the expression of four significantly altered genes from differential gene expression (DEG) analysis of DG neurons on the pseudotime embedding, the color scale of the raw counts was adjusted by log10 transformation. **g,** Spatial cell map colored by pseudotime for DG region. Scale bar, 100 µm. **h,** p-Tau signal quantification around plaques. p-Tau+ pixels (intensity > threshold, see Method) were quantified at different distance intervals (0-10, 10-20, 20-30, 30-40, and 40-50 µm) to the Aβ plaque in the cortex and subcortical regions of the TauPS2APP 13 month sample. Y-axis values were normalized by the total p-Tau signal of each brain region. **i,** Stacked bar chart showing the composition of p-Tau positive excitatory neurons and inhibitory neurons in each AD sample defined by the ratio of tau positive pixels to the area of each cell (see Methods). **j,** Synaptic gene ontology term enrichment of DEGs (p-value < 0.05) identified from the p-Tau positive CA1 neurons versus p-Tau negative CA1 neurons (Methods) using SynGO. Color of the sunburst plot represents enrichment -log10 Q-value at 1% FDR.

We investigated neuron subtype compositions and their transcriptomic profiles in relation to Aβ plaques. In the cortex, there was a paucity (strong relative reduction) of all types of neurons adjacent to plaques (<10 µm, Fig. 6c,d and Extended Data Fig. 6e,f), anti-correlating with the large increase in density of microglia close to plaque. In the hippocampus the impact of plaque on the density of neuron subtypes was difficult to interpret because of the anatomic organization of the hippocampus and because amyloid plaques were concentrated in the neuropil, relatively far from the cell body layers, and mostly in the molecular layer of dentate gyrus (Fig. 6c and Extended Data Fig. 6e).

Pseudotime trajectory analysis of DG neuron transcriptomes showed a bifurcate trajectory (Fig. 6e,f): At 8 months, when there were very few plaques near DG (1 plaque within the 50 µm distance to DG), the pseudotime distribution of DG cells in TauPSAPP and control mice were largely indistinguishable (Fig. 6e,g). At 13 months, accompanied by the increased number of plaques (13 plaques) in DG, the pseudotime trajectory diverged into two branches corresponding to the DG populations in TauPS2APP (lower left branch) and control mice (lower right branch) (Fig. 6e). DEG analyses on pseudotime trajectory embedding further revealed that *Ccnb2*, *Dapk1*, and *Tnik* were upregulated in the TauPS2APP DG neurons versus control, while *Ngrn* was downregulated (Fig. 6f, Supplementary Table 3). *Dapk1* is involved in neuronal death regulation whereas *Tnik* has been implicated in dentate gyrus neurogenesis^46^, so these results may indicate altered neurogenesis in DG in TauPS2APP mice, which is related to a recent report showing that the adult hippocampal neurogenesis activity in DG sharply declines in human patients of AD^47^.

To investigate the neuronal transcriptomic alterations induced by tauopathy, we first quantified the ratio of p-Tau positive pixels to the total pixel area of neuronal cell bodies and defined p-Tau positive neurons as those in which this ratio is greater than 7% (see Methods). In TauPS2APP mice, there were 4 times more p-Tau positive neurons at 13 months than 8 months. At 8 months the majority of p-Tau positive neurons were CTX-Ex2 excitatory neurons, whereas at 13 months the majority of p-Tau positive neurons were the CA1 excitatory neurons. Inhibitory neurons account for <20% p-Tau positive neurons: most of them were *Pvalb* neurons at 8 months whereas at 13 months most of them came from the *Sst* population (Fig. 6i). We also quantified the p-Tau signal around the plaques and found that the p-Tau signal was enriched within 10 µm distance near plaques (Fig. 6h). Considering neuronal cell bodies were relatively depleted within the 10 µm range, the observed p-Tau signals likely correspond to dystrophic (injured) neurites^35^.

We finally examined the DEGs of all neuronal types in response to Aβ plaques and p-Tau. GO term analysis of general neuronal DEGs (TauPS2APP versus control) revealed upregulation of protein kinase activity, downregulation of protein transport and section, and downregulation of mRNA processing (Extended Data Fig. 6g). The DEGs of the dentate gyrus region were related to downregulation of brain development (Extended Data Fig. 6h). SynGO analysis^36^ of DEGs in p-Tau positive CA1 neurons at 13 months are clustered in synaptic vesicle cycle, synaptic vesicle endocytosis and postsynapse organization (Fig. 6j). Neuron-type resolved DEG analysis (Supplementary Table 3) revealed that *Ccnb2* (G2/mitotic-specific cyclin-B2) was identified among top upregulated DEGs from all neuronal types in TauPS2APP models, as well as genes involved in cell death (*i.e. Ddit3, Dapk*), suggesting a reactivation of the cell cycle and supporting the notion that neuronal reentry into the cell cycle is a mediator of neurodegeneration and cell death^48, 49^.

### Integrative analysis of disease-associated cells and genes in AD pathology

The analyses above focused on dissecting disease-associated subtypes and DEGs within major brain cell types. In order to synthesize a comprehensive picture of AD gene pathways from multiple cell types, we performed cell-type-resolved Gene Set Enrichment Analysis (GSEA)^50, 51^ using DEGs from four major cell types (microglia, astrocytes, oligodendrocytes, and neurons) comparing TauPS2APP versus control. The results (Fig. 7a,b) showed that most of the significantly enriched terms of upregulated genes in non-neuronal cell DEGs were in the biological processes of inflammatory response, glial cell differentiation, gliosis, cytokine production, and regulation of neuronal death, while those of neurons involve synaptic signaling and protein kinase activity. Neurons in the TauPS2APP mice also showed significant downregulation of RNA processing/splicing and protein transport (Fig. 7a,b).

**Fig. 7.**
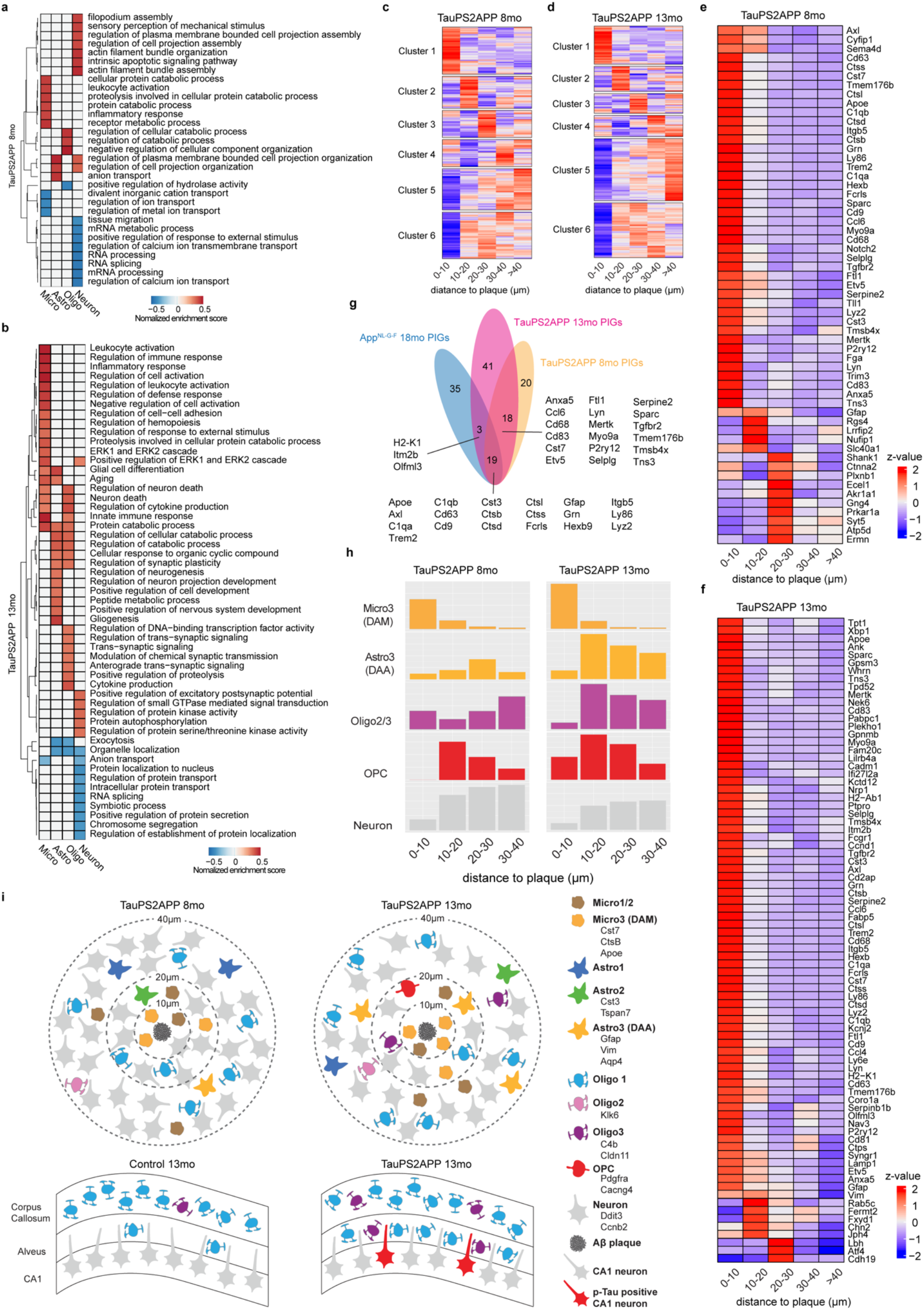
Integrative spatiotemporal analysis of disease-associated cell types and gene programs. **a,b,** GSEA heatmap showing the significant (nominal p-value < 0.05) biological process related terms enriched in DEGs of each interested cell type in TauPS2APP samples at 8 months (**a**) and 13 months (**b**). Terms are filtered by term size: 20-1000, Color of the tiles represents normalized enrichment score. **c,d,** Matrix plot showing the gene clustering results in each distance interval (0-10, 10-20, 20-30, 30-40, >40 µm) around the Aβ plaque in TauPS2APP 8-month sample (**c**) and TauPS2APP 13-month sample (**d**). Colored by row-wise z-score. **e,f,** Matrix plot showing the Plaque Induced Genes (Enriched in 0-40 interval, adjusted p-value < 0.01) in each distance interval (0-10, 10-20, 20-30, 30-40, >40 µm) around the Aβ plaque in TauPS2APP 8-month sample (**e**) and TauPS2APP 13-month sample (**f**). Colored by row-wise z-score. **g,** Venn Diagram highlighting the overlap of SDEGs in the TauPS2APP 8- and13-month samples with SDEGs in TauPS2APP and previously reported PIGs in 18 month *App_NL-G-F_* mice^33^. **h,** Spatial histograms of Micro3 (DAM), Astro3 (DAA), Oligo2/3, OPC and neuronal cells around Aβ plaque in the TauPS2APP 8-month sample (left) and 13-month sample (right). **i,** Schematic diagram showing the spatial distribution of different cell types around Aβ plaque and oligodendrocyte subtypes in hippocampal alveus in the TauPS2APP mouse model. The number of cells in the schematic diagram represents the approximate ratio of cell number for each cell type.

Besides the DEGs defined by comparison of TauPS2APP vs. control samples, we also identified plaque-associated *spatial* DEGs (SDEGs) by clustering genes based on their expression profiles as a function of distance from amyloid plaques (Fig. 7c,d). We identified six clusters of SDEGs: Clusters 1-3 are specifically upregulated at different distances near plaques, and can be regarded as plaque-induced genes (PIGs, listed in Fig. 7e,f); Cluster 4-6 are relatively downregulated near plaques and enriched with neuronal genes (Supplementary Table 4). We identified 57 PIGs at 8 months and 81 PIGs in the TauPS2APP sample at 13 months (Fig. 7g). 37 out of 57 PIGs in the 8-month TauPS2APP sample were also found in the 13-month samples (Fig. 7g). The PIGs enriched within 10 µm distance from plaques were mainly DAM marker genes, such as *Trem2*, *Cst7*, *Ctsb*, *Apoe* and *Cd9* (Fig. 7e,f). The DAA marker *Vim* was upregulated in the regions 10 to 20 µm away from plaques in AD 13-month samples (Fig. 7f), which is consistent with our previous finding that DAA enriched in the 10-20 µm ring (Fig. 4). Although SDEGs in Cluster 4-6 (Supplementary Table 4) are down-regulated at the 0-30 µm distance from plaques and not defined as PIGs, they are also valuable to reveal potential disease mechanisms. For example, an interesting set of SDEGs was induced at 8 months with peak expression at 30-40 µm from plaque, including *Elavl4*, an RNA binding protein that is implicated in early dysregulation of splicing in frontotemporal dementia^52^. Comparing our PIGs with previously reported PIGs identified from 18-month AppNL-G-F mice^33^, 22 out of 81 and 19 out of 57 PIGs in our AD 8-month and 13-month samples, respectively, overlapped with previously reported PIGs (Fig. 7g).

To validate the findings from the study, we repeated the STARmap PLUS experiment on another set of TauPS2APP mice, this time using a focused subset of genes (64 key cell-type marker genes and disease-associated genes). This focused 64 gene analysis yielded cell clustering results, DEG data and spatial map information that was highly consistent with the major conclusions derived from the 2766-gene datasets (Supplementary Tables 1,2, Extended Data Fig. 7) including the spatial enrichment of various glial cells surrounding plaques (Extended Data Fig. 7d), the infiltration of oligodendrocytes in hippocampus alveus (Extended Data Fig. 7e), as well as cell-type and spatially resolved DEGs (Extended Data Fig. 7f-h).

## Discussion

Here we developed STARmap PLUS for *in situ* detection of RNAs and proteins in the same tissue section at subcellular resolution. Compared to STARmap, we have substantially improved the number of *in situ* sequenced genes and enabled a simultaneous profiling of RNAs and proteins in intact hydrogel-tissue scaffolds. This method development enables a multimodal analysis of spatially expressed RNAs and proteins, bringing an opportunity to study biological systems in a more comprehensive manner. This method is readily applicable to a variety of samples for study of physiological and pathological mechanisms in healthy and diseased tissues by integrative mapping of single-cell transcriptomic states, tissue morphology and disease markers.

### Analysis framework

We developed STARmap PLUS to enable high-resolution spatial transcriptomics in combination with microscopic protein localization. In a mouse model of AD exhibiting amyloidosis, tauopathy and neurodegeneration we applied this high-resolution and multimodal *in situ* mapping with four computational analysis strategies to identify disease-associated cell populations and gene programs: (1) hierarchical cell clustering analysis to pinpoint disease-associated cell subtypes, (2) pseudotime trajectory analysis to reconstruct the transitions of cell states during disease progression, (3) spatial analysis to recognize changes in cell types and cell states in physical proximity of Aβ plaques and tauopathy; (4) differential gene expression analysis in spatiotemporal relationship with amyloid and p-Tau and disease stage to identify disease-associated gene pathways. This analysis framework successfully traced how AD hallmark pathologies correlate with gene pathways that drive inflammation, gliosis, and neurodegeneration across different cell types.

### Core-shell structures surrounding plaques and cascades of cell-state transitions

Applying our methods to the TauPS2APP mouse AD model at two different ages, we have constructed a cell-type and cell-state resolved spatiotemporal map of the TauPS2APP mouse model (schematized in Fig. 7i). Specifically, in response to the early emergence of Aβ plaques (at 8 months or earlier), microglia are the primary responders, closely aggregating around plaques (<10 µm distance away from plaques, Fig. 7i). Integrative spatial and pseudotime analysis of microglia subtypes suggested a cell state transition from Micro1 and Micro2 to Micro3 subtype (DAM) associated with microglia cells accumulating around Aβ plaques (Fig. 3 and Extended Data Fig. 3). The transition from Astro2 to Astro3 (DAA) in the layer (10-20 µm) next to the DAM microglia was seen at a later stage (13 months), which is consistent with an induction of DAA by DAM. Indeed, a previous study showed that reactive astrocytes with the high expression of DAA marker genes *Gfap* and *Vim*, were induced by activated microglia^21^. Like DAA, Oligo2/3 accumulation near plaque (10-30 µm) occurred later (13 months) (Fig. 7h). Interestingly, we discovered that OPCs were enriched in the regions 10-30µm away from plaques at both 8-month and 13-month stages, indicating potential *in situ* proliferation and differentiation of OPCs to oligodendrocytes. In contrast to glia, neuron density around plaque declined from 8 months to 13 months (Fig. 7h,i).

Based on the collective data, we propose a core-shell structure of glial cells surrounding Aβ plaques where the DAM emerge early in disease near plaques as the core, and the shell is a gliosis zone enriched for DAA, OPC, and oligodendrocytes that develops at a later disease stage, perhaps dependent on formation of the inner ring of reactive microglia. Because STARmap PLUS only captures snapshots from different disease stages, the dynamic cell-type and state transitions inferred from pseudotime trajectory reconstruction and spatial patterns need future verification by live-cell imaging or *in vivo* cell-fate tracing approaches.

### Oligodendrocytes and tauopathy

In the TauPS2APP model, hyperphosphorylated tau (as detected with AT8 antibody) was mainly found in CA1 excitatory neuronal bodies and its upper layers of axon tracts (alveus), where it is strongly associated with the infiltration of Oligo1 and Oligo3 subtypes regardless of the presence or absence of plaques (schematized in Fig. 7i). It is unclear whether the infiltration of Oligo1 and Oligo3 is a reactive mechanism to support tauopathy-injured axons, repair damaged myelin, or whether Oligo1 and Oligo3 instead exacerbate tauopathy. It is possible that Oligo1 and Oligo3 might feature distinct responses and exert different impacts, for example, Oligo 1 being homeostatic and protective while Oligo 3 is detrimental in response to tau-driven changes. The reciprocal functional interactions between oligodendrocytes and neurons (beyond oligodendrocytes providing electrical insulation and metabolic support for axons) are increasingly appreciated and studied^53^. Single-cell RNAseq studies of human AD brain tissues have noted major transcriptomic changes in oligodendrocytes, but without spatial information in relation to plaque and p-Tau^12^. Enabled by high-resolution spatial transcriptomic analysis of oligodendrocytes in the TauPS2APP mice, we identified oligodendrocytes subtypes that are associated with tauopathy. Further pathway analysis implicated oligodendrocyte DEGs in the regulation of neuron death, cytokine production, synaptic plasticity, and anterograde synaptic signaling (Extended Data Fig. 5h), pointing to a potential link between inflammation and axonal tauopathy mediated by oligodendrocytes.

### Implications for neurodegeneration mechanisms in AD

The overall density of neuronal cells in cortex declines near Aβ plaques (Fig. 6c,d), implying a loss of neurons due directly to Aβ toxicity or indirectly via the effects of amyloid on microglia and other glia surrounding the plaque. This notion is supported by the accumulation of p-Tau in dystrophic neurites near to plaque. We also cannot exclude that neurons are merely displaced from plaque-adjacent regions by reactive glia; however, we note that the TauPS2APP mice show macroscopic brain volume loss by 13 months of age^35^. Consistent with neuronal loss around plaques, we note that neurons in DG, where a large amount of Aβ plaques appeared by 13 months, showed altered transcriptional profile in intracellular protein transport, RNA processing and cell cycle pathways related to cell stress and neurodegeneration as well as molecular pathways related to synapse structure and organization. Meanwhile, all neuronal types examined in cortex and hippocampus of TauPS2APP share some common DEGs (i.e. *Ccnb2, Ddit3*), perhaps suggesting a general mechanism of cell cycle reactivation in neurodegeneration. The core-shell glial structure around plaques implies that microglia may cause neurodegeneration in part through their crosstalk with astrocytes^21^ and oligodendrocytes. Different subtypes of the oligodendrocyte lineage respond differently to plaques and tauopathy, suggesting distinct modes of oligodendrocyte recruitments (*e.g.* microglia-oligodendrocyte/OPC interactions near plaques, neuron-oligodendrocyte interactions in tauopathy). Future studies in human patient samples and other AD disease models are needed to test the potential pathogenic mechanisms revealed by STARmap PLUS analysis of the AD brain.

## Acknowledgements

We thank Zefang Tang, Hailing Shi, Ming Pan, Qiang Li and Zuwan Lin and Sarah Wade for technical assistance, Han Xu (Peking University) and Haowei Meng (Peking University) for helpful discussions and thoughtful comments on the manuscript. X.W. acknowledges the support from the Searle Scholars Program, Thomas D. and Virginia W. Cabot Professorship, and Edward Scolnick Professorship.

## Author contributions

H.Zeng and J.R. optimized STARmap PLUS probe design. H.Zeng and Y.Z. performed STARmap PLUS experiments. J.H. and H.Zhou performed computational analyses. W.L.M. prepared animal samples. H.Zeng, J.H., H.Zhou, and X.W. analyzed and interpreted the data with the inputs from W.J.M., B.D., C.J.B., S.-H.L, and M.S. All authors contributed to writing and revising the manuscript and approved the final version. X.W. and M.S. conceptualized and supervised the project.

## Methods

### Mice

All animal procedures followed animal care guidelines approved by the Genentech Institutional Animal Care and Use Committee (IACUC) and animal experiments were conducted in compliance with IACUC policies and NIH guidelines. The mice used for STARmap PLUS include the pR5-183 line expressing the P301L mutant of human tau and PS2_N141I_ and APP_swe_ (PS2APP^homo^; P301L^hemi^) and non-transgenic control.

### Tissue collection and sample preparation for STARmap PLUS

Animals were anesthetized with isoflurane and rapidly decapitated. Brain tissue was removed, placed in O.C.T, then frozen in liquid nitrogen and kept at -80 °C. For the tissue sectioning, mouse brains were transferred to cryostat (Leica CM1950) and cut into 20 µm thick slices in coronal sections at -20 °C. The slices were attached to each well of glass-bottom 12-well plates pre-treated by methacryloxypropyltrimethoxysilane (Bind-Silane) and poly-D-lysine (PDL). The brain slices were fixed with 4% PFA in 1X PBS buffer at room temperature for 15 min, then permeabilized with -20 °C methanol and placed at -80 °C for an hour before hybridization.

### STARmap PLUS to detect spatial RNA and protein signals

The samples were taken from -80 °C to room temperature for 5 min and then washed with PBSTR buffer (0.1% Tween-20, 0.1 U/µl SUPERase·In RNase Inhibitor in PBS). After washing, the samples were incubated with 300 µl of 1X hybridization buffer (2X SSC, 10% formamide, 1% Tween-20, 0.1 mg/ml yeast tRNA, 20 mM Ribonucleoside vanadyl complexes, 0.1 U/µl SUPERase·In RNase Inhibitor and pooled SNAIL probes at 1 nM per oligo) in a 40 °C humidified oven with shaking and parafilm wrapping for 36 h. The samples were washed by PBSTR twice and high-salt washing buffer (4X SSC dissolved in PBSTR) once at 37 °C. Finally, the samples were rinsed with PBSTR once at room temperature. The samples were then incubated with a ligation mixture (1: 10 dilution of T4 DNA ligase in 1X T4 DNA ligase buffer supplemented with 0.5 mg/ml BSA and 0.2 U/µl of SUPERase·In RNase inhibitor) at room temperature for two hours with gentle shaking. After ligation, the samples were washed twice with PBASR buffer and then incubated with rolling circle amplification (RCA) mixture (1: 10 dilution of Phi29 DNA polymerase in 1X Phi29 buffer supplemented with 250 µM dNTP, 20 µM 5-(3-aminoallyl)-dUTP, 0.5 mg/ml BSA and 0.2 U/µl of SUPERase·In RNase inhibitor) at 30 °C for two hours with gentle shaking. Subsequently, the samples were washed twice with PBST (0.1% Tween-20 in PBS) and blocked with blocking solution (5 mg/ml BSA in PBST) at room temperature for 30 min. The samples were then incubated with Phospho-Tau (Ser202, Thr205) Antibody (Thermo, MN1020B, 1:100 dilution in blocking solution) for 2 hours at room temperature. The samples were washed with PBST three times for 5 min each. Next, the samples were treated with 20 mM Acrylic acid NHS ester in PBST for 1 hour and rinsed once with PBST. The samples were incubated in the monomer buffer (4% acrylamide, 0.2% bis-acrylamide in 2X SSC) for 15 min at room temperature. Then the buffer was aspirated, and a 35 µl polymerization mixture (0.2% ammonium persulfate, 0.2% tetramethylethylenediamine dissolved in monomer buffer) was added to the center of the sample and immediately covered by Gel Slick-coated coverslip. The polymerization reaction was undergone for 1 hour at room temperature (N_2_) and washed by PBST twice for 5 min each. Subsequently, the samples were treated with dephosphorylation mixture (1:100 dilution of Shrimp Alkaline Phosphatase in 1X CutSmart buffer supplemented with 0.5 mg/ml BSA) at 37 °C for 1 hour and washed by PBST three times for 5 min each.

For SEDAL sequencing, each cycle began with treating the sample with stripping buffer (60% formamide and 0.1% Triton-X-100 in H_2_O) at room temperature for 10 min twice, followed by PBST washing for three times, 5 min each. The sample was incubated with a sequencing mixture (1: 25 dilution of T4 DNA ligase in 1X T4 DNA ligase buffer supplemented with 0.5 mg/ml BSA, 10 µM reading probe, and 5 µM fluorescent oligos) at room temperature for at least 3 hours. The samples were washed by washing and imaging buffer (10% formamide in 2X SSC) three times, 10 min each, then immersed in washing and imaging buffer for imaging. Images were acquired using Leica TCS SP8 confocal microscopy. Eight cycles of imaging were performed to detect 2,766 genes.

After 8-round *in situ* sequencing for 2766-gene samples and 4-round *in situ* sequencing for 64-gene samples, the sample was incubated in X-34 solution (10 µM X-34, 40% ethanol and 0.02 M NaOH in 1X PBS) at room temperature for 10 min, followed by quick washing with 1X PBS for 3 times. The samples were incubated 80% EtOH for 1 min and then washed with PBS 3 times, 1 min each. Then the samples were incubated with the Goat anti-Mouse IgG (H+L) Cross-Adsorbed Secondary Antibody, Alexa Fluor 488 (Thermo, A-11001, 1:80 dilution in blocking solution) at room temperature for 12 h. The sample was washed three times with PBST for 5 min each. The samples were incubated with the 500nM 19-nt fluorescent oligo complementary to DNA amplicon in PBST at room temperature for 1h, then washed by PBST three times for 5 min each. Propidium Iodide (PI) staining was performed following the manufacturer’s instruction for the purpose of cell segmentation. Another round of imaging was performed to detect spatial protein signals.

### STARmap PLUS Image Processing

All of the image processing steps were implemented using MATLAB R2019b and related open-source packages in Python 3.6 and applied according to^32^.

#### Image Preprocessing

A multi-dimensional histogram matching was performed on each tile with MATLAB function ‘imhistmatchn ’. It used the image from the first color channel in the first sequencing round as a reference to uniform the illuminance and contrast level of all other images in the current imaging position. Additionally, a customized tophat filtering was applied to the sequencing images to further enhance the signal and suppress the background noise.

#### Image Registration

Image registration was applied according to^32^. Global image registration was accomplished using a three-dimensional fast Fourier transform (FFT) to compute the cross-correlation between two image volumes at all translational offsets. The position of the maximal correlation coefficient was identified and used to translate image volumes to compensate for the offset. Then a non-rigid registration was applied with MATLAB function ‘imregdemons ’ to further align images in different sequencing rounds.

#### Spot Calling

After registration, individual dots were identified separately in each color channel on the first round of sequencing. For this experiment, amplicon dots were identified by finding local maxima in 3D with MATLAB function ‘imregionalmax ’. Dots with intensity at their centroids less than the threshold were removed. Because the dot was approximately 6 pixels in diameter on the xy plane, the dominant color for that dot across all four channels on each round was determined by a 5×5×3 voxel volume surrounding the dot centroid. The integrated intensity of the voxel volume in each channel was used for color determination. In this case, each dot in each round had a L2 normalized vector with four elements. The color of each dot was determined by the corresponding channel with the highest value in the vector. Dots with multiple maximum values in the vector were discarded.

#### Barcode Filtering

Dots were first filtered based on quality scores (average of -log(color vector value in dominant channel) across all sequencing rounds). The quality score quantified the extent to which each dot on each sequencing round came from one color rather than a mixture of colors. The barcode codebook was converted into color space based on the expected color sequence following the 2-base encoding of the barcode DNA sequence. Dot color sequences that passed the quality threshold and matched sequences in the codebook were kept and identified with the specific gene that that barcode represented; all other dots were rejected. The high-quality dots and associated gene identities in the codebook were then saved for downstream analysis.

#### 2D Cell Segmentation

Nuclei were automatically identified by applying a pre-trained 2D machine learning model from the StarDist package^54^ to a maximum intensity projection of the stitched DAPI channel following the final round of sequencing. Then the cell locations were extracted from the segmented DAPI image and used as markers for cell body segmentation. Cell bodies were represented by an overlay of stitched Nissl staining and merged amplicon images. Finally, a marker-based watershed transform was applied to segment the thresholded cell bodies. Points overlapping each segmented cell region in 2D were then assigned to that cell, to compute a per-cell gene expression matrix.

#### Protein image preprocessing

p-Tau images were processed using a customized Fiji macro^55^. A rolling ball background subtraction with a radius equal to 5 was applied to each image. Then each image was processed with a Gaussian filter with a sigma value equal to 2 and followed by a maximum entropy binarization.

#### Cell type classification

A two-level clustering strategy was applied to identify both major and sub-level cell types in the dataset. Processing steps in this section were implemented using Scanpy v1.4.6^56^ and other customized scripts in Python 3.6 and applied according to Wang et al (2018)^32^. After filtration, normalization, and scaling, principal component analysis (PCA) was applied to reduce the dimensionality of the cellular expression matrix. Based on the explained variance ratio, the top PCs were used to compute the neighborhood graph of observations. Then the Leiden algorithm was used to identify well-connected cells as clusters in a low dimensional representation of the transcriptomics profile. The cells were displayed using the Uniform Manifold Approximation and Projection (UMAP) and color-coded according to their cell types. The cells for each interested top-level cluster were then extracted and sub clustered using the same workflow again.

### Pseudotime and Trajectory Analysis

An R package Monocle3^44^ (https://cole-trapnell-lab.github.io/monocle3/) was utilized for pseudotime calculation. Monocle3 firstly learned a principal graph (via learn_graph() function) in the three-dimensional UMAP space using a dimensionality-reduced representation of original data. In detail, Monocle3 ran a k-means clustering algorithm and selected “landmark cells” by first mapping cells to their nearest k-means points, then choosing the cell with highest local density as landmark cell. A principal graph was learned within these landmark cells rather than the whole dataset to reduce running time using simplePPT algorithm^57^! After generating the principal graph, the graph was refined by initially performing a depth-first visitation of the nodes in the graph. For nodes with degree (the number of edges connected to this node) ≤ 2, no operation was performed. For nodes with degree > 2, Monocle3 computed the diameter path for each subtree of the node not visited, and removed the subtree if its path length is less than a dedicated length specified by the user.

To compute pseudotime from the principal graph, firstly all cells were mapped to their nearest principal points based on Euclidean distance in the UMAP embedding. Next, for each edge in the graph, all cells mapped to the endpoints were orthogonally projected to the nearest points on the graph edge. Based on the projected position of cells, suppose the endpoints are ***a*** and ***b*** and here we have cells named ***c_i_ , c_j_*** , with projection ***p(c_i_ ) , p(c_j_ )*** . We can order the cell projections on the edge and suppose the order is ***a < p(c_i_ ) < p(c_j_ ) < b***. Then edge ***(a,c_i_ )*** and ***(b,c_j_ )*** will be added to the final graph. After specifying the root node(s) by user, the pseudotime for each cell can be assigned by calculating the geodesic distance from root node to the cell node. In this case, raw expression matrix of each cell type: Microglia, Astrocyte, Oligodendrocyte/Oligodendrocyte precursor cell (OPC) and Dentate gyrus (DG), was used separately as the input of Monocle3. To visualize the results, we used the Monocle3-generated UMAP embedding with palettes representing normalized pseudotime (rescaled to 0 ∼ 1), cell types, and sample identities.

### Spatial Analysis

#### Plaque Segmentation

The plaques were segmented from the thresholded image of plaque channel by using the ‘bwlabel ’ function in EBImage package^58^. Then, the size and the center of each plaque were calculated by using ‘computeFeatures.moment ’ and ‘computeFeatures.shapè functions respectively. Finally, plaques with an area greater than 400 pixels (∼ 40 µm^2^) were kept for the downstream analysis.

#### Cell Composition around Plaques

As filtered plaques were acquired in the last step, we dilated the plaque images 5 times with steps of 10 microns. Next, we counted the number of cells for every cell type that fall into different intervals. The ranges were set from 0-10µm (Ring 1) to 40-50µm (Ring 5). The statistics were normalized by calculating the percentage and density of each cell type in a ring. The graphical explanation of this analysis is shown in Fig. 2d. For the overall statistics, we calculated the percentage of each cell type in the whole sample. To test the significance of cell enrichment in both density and proportion aspects in specific intervals, two kinds of statistical tests: one sample t-test and chi-square test (Fisher’s Exact test) were utilized. One sample t-test was used to compare the density values for each cell type in one interval around every plaque with the average density. According to the difference of mean value of density sequence and the average density, an alternative hypothesis was set correspondingly to test whether the density was significantly higher or lower. For the chi-square test, raw cell counts in each interval were used for the test.

### Differential Expression Analysis

Before performing DE analysis, the dataset was normalized according to the following steps: 1. Divide the gene counts in a sample by the median of *total counts per cell* for that sample and multiply by the scale factor, which was defined as the mean value of median of *total counts per cell* for all samples; 2. Perform log2 transformation by adding a pseudo-count of one. DE genes were identified by performing Wilcoxon-Rank Sum test between two groups of cells using the ‘FindMarkers ’ function in Seurat. For ‘disease vs. control’ comparison of specific cell types, the two groups of cells were extracted from TauPS2APP and control samples and compared. As for the comparison between cells close to plaque versus cells far away from plaque, the cell was defined as ‘near plaque’ if its distance to the nearest plaque was less than 25µm and all other cells with distance to plaque greater than 25µm were defined as ’away from plaque’. In the comparison of ‘CA1 Tau+ vs Tau-’, Tau+ CA1 cells were defined according to the fraction of Tau signal area to the cell body’s area. The threshold was set to 0.07.

In order to filter out lowly expressed genes, genes that were expressed by less than 10% of cells in either group of the comparison were excluded. We also applied the following threshold values on the generated gene list to filter out non-significant genes: absolute value of log fold change > 0.1, p-value < 0.05.

To visualize the DE result, we used the ‘EnhancedVolcanò package (https://github.com/kevinblighe/EnhancedVolcano) to generate the volcano plot. DE genes with logFC > 0 were colored in red while others were colored in blue. For those significant genes (p-value < 0.05) but failed to pass the LogFC threshold were green tinted. All other non-significant genes were colored in gray. Note, some genes with extremely high -log(P-value) or logFC were capped.

### Gene Set Enrichment Analysis

Gene Set Enrichment Analysis (GSEA v4.1.0)^50, 51^ was used to perform pathway analysis for each cell type with a LogFC ranked gene list whose genes were detected in a minimum fraction of 5% cells in the targeted cell type. Gene Ontology Biological Processes (GO:BP) database was selected as the gene sets database to perform GSEA. To limit the size of gene sets subjected to enrichment analysis, the minimum and maximum size of gene sets were set to 20 and 1000. Number of permutations was set to 1000. Results were exported in tab-delimited format to be used as input for further functional enrichment analysis. The SynGO enrichment tool^36^ was used to further characterize the synapse functions enriched in DEGs from excitatory neurons, inhibitory neurons, and CA1 cells with Tau pathology. Brain expressed genes were used as the background gene list.

### Grid Analysis

To analyze the scattered p-Tau signal, we firstly performed rasterization on p-Tau images with resolution at 20µm (See heatmap Extended Data Fig. 5f). For each block, we counted the number of cells for each major cell type and the p-Tau signal intensity. Based on the distribution of p-Tau intensity and the existence of plaque, blocks were classified into 4 groups: No p-Tau, Low p-Tau (intensity of p-Tau in 1 to 50% range of population with non-zero p-Tau signal), High p-Tau w/ plaque (51-100% range with plaque) and High p-Tau w/o plaque (51-100% range without plaque). The significance of cell type enrichment was tested using one-sample t-test comparing the density values of each group with average density.

### Plaque Induced Genes

To identify potential gene expression patterns around plaque, we firstly calculated the average expression value for each gene in 5 intervals: 0-10µm, 10-20µm, 20-30µm, 30-40µm, >40µm. The averaged expression matrix was calculated separately for disease samples in 8 months and 13 months. Next, K-means clustering was applied on the matrix respectively and 6 clusters were obtained in each sample. We tested whether the averaged expression level of the cells within the high expression interval was significantly higher than that of other cells outside the interval using Wilcoxon signed-rank test (With p-value < 0.05, Supplementary Table 4). Plaque induced genes (PIGs) were defined as significantly enriched genes within the 0-30 µm distance surrounding plaques (Cluster 1-3). ‘ComplexHeatmap’ package^59^ is utilized for heat map visualization.

### Quantification and Statistical analysis

The statistical tests and number of independent replicates per experiment are indicated in the Figure legends. The statistical significance from STARmap PLUS sequencing experiments are detailed in the Method details section.

One-sample t-test was used to test the significance of cell type enrichment by comparing the density values of each cell type in each interval with average density.

### Data availability

The STARmap PLUS sequencing data are available on Single Cell Portal (https://singlecell.broadinstitute.org/single_cell/study/SCP1375).

### Code availability

The code for the STARmap PLUS image analysis is available on Zenodo (https://zenodo.org/record/5842625#.YeAvYljML0o)

---

**Extended Data Fig. 1.**
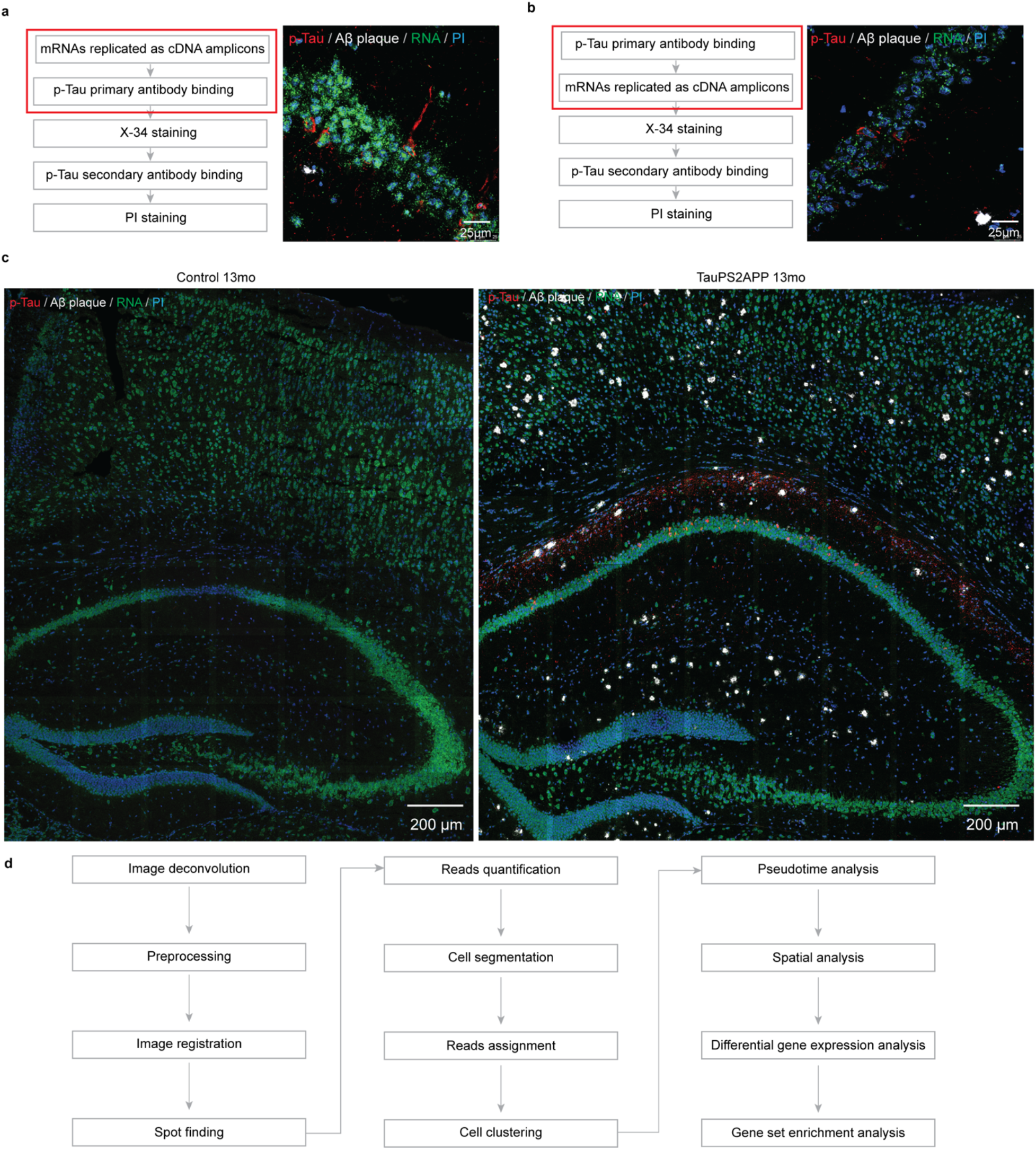
Development of the STARmap PLUS method. **a,** STARmap PLUS procedure where p-Tau primary antibody staining was performed after mRNA *in situ* hybridization and amplification. The imaging results showed strong signals from both cDNA amplicons and proteins. **b,** Alternative procedure where the p-Tau primary antibody staining was conducted before mRNA *in situ* hybridization and amplification. The imaging results showed much weaker signal from cDNA amplicons, suggesting RNA degradation during antibody incubation and washing steps. PI staining, propidium iodide staining of cell nuclei. **c,** The ninth round of the imaging detected cell nuclei, cDNA amplicons, and protein signals in the 13-month control mouse brain (left) and TauPS2APP mouse brain (right). Blue, Propidium Iodide (PI) staining of cell nuclei. Green, fluorescent DNA probe staining of all cDNA amplicons. White, X-34 staining of Amyloid β plaque. Red, immunofluorescent staining of p-Tau (AT8 primary antibody followed by fluorescent goat anti-mouse secondary antibody). **d,** The flowchart of the STARmap PLUS data analysis pipeline.

**Extended Data Fig. 2.**
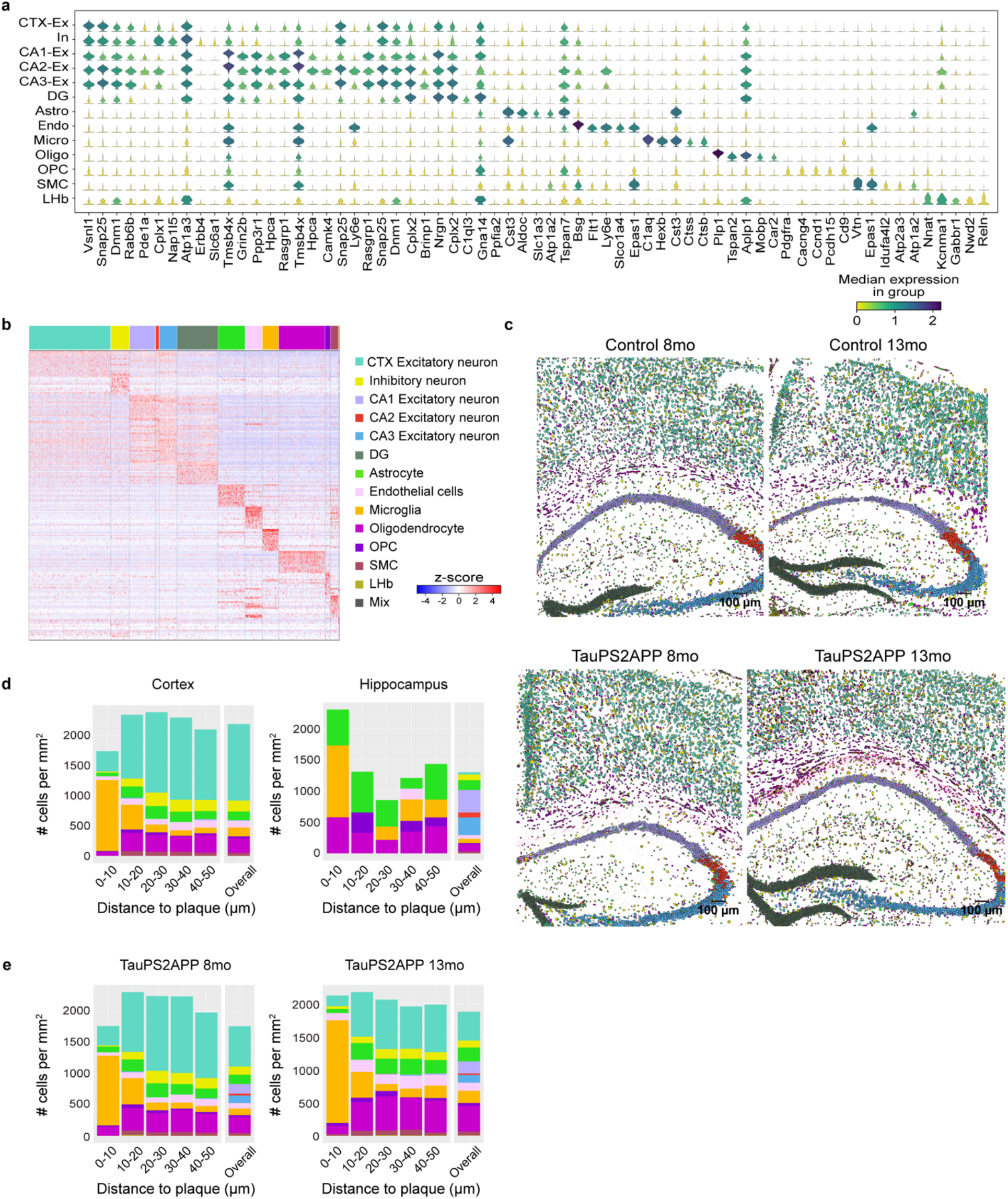
Top-level cell type classification. **a,** Stacked violin plot for representative gene markers aligned with each top-level cell type of 2,766 genes dataset. **b,** Gene expression heatmaps for representative markers aligned with each top-level cell type of 2,766-gene datasets. Expression for each gene is z-scored across all genes in each cell. **c,** Spatial atlas of top-level cell types in cortex and hippocampus regions of 4 samples in the 2,766-gene dataset. Scale bars, 100 µm. **d,** Representative spatial distribution of cell type compositions around Aβ plaque for TauPS2APP 8-month sample in both cortex and hippocampus. Stacked bar plot showing the density (cell count per mm^2^) of each major cell type at different distance intervals (0-10, 10-20, 20-30, 30-40, 40-50 µm) to the Aβ plaque. The cell density of each major cell type in each area was included as the reference for comparison. **e,** Cell-type composition around Aβ plaque at different distance intervals in both 8- and 13-month samples of the 2,766-gene dataset. Stacked bar plot showing the averaged density (count per mm^2^) of major cell types in the cortex and hippocampus from different distance intervals (0-10, 10-20, 20-30, 30-40, 40-50 µm) around the Aβ plaque. The overall cell density of each cell type in each region was included as the reference for comparison(overall).

**Extended Data Fig. 3.**
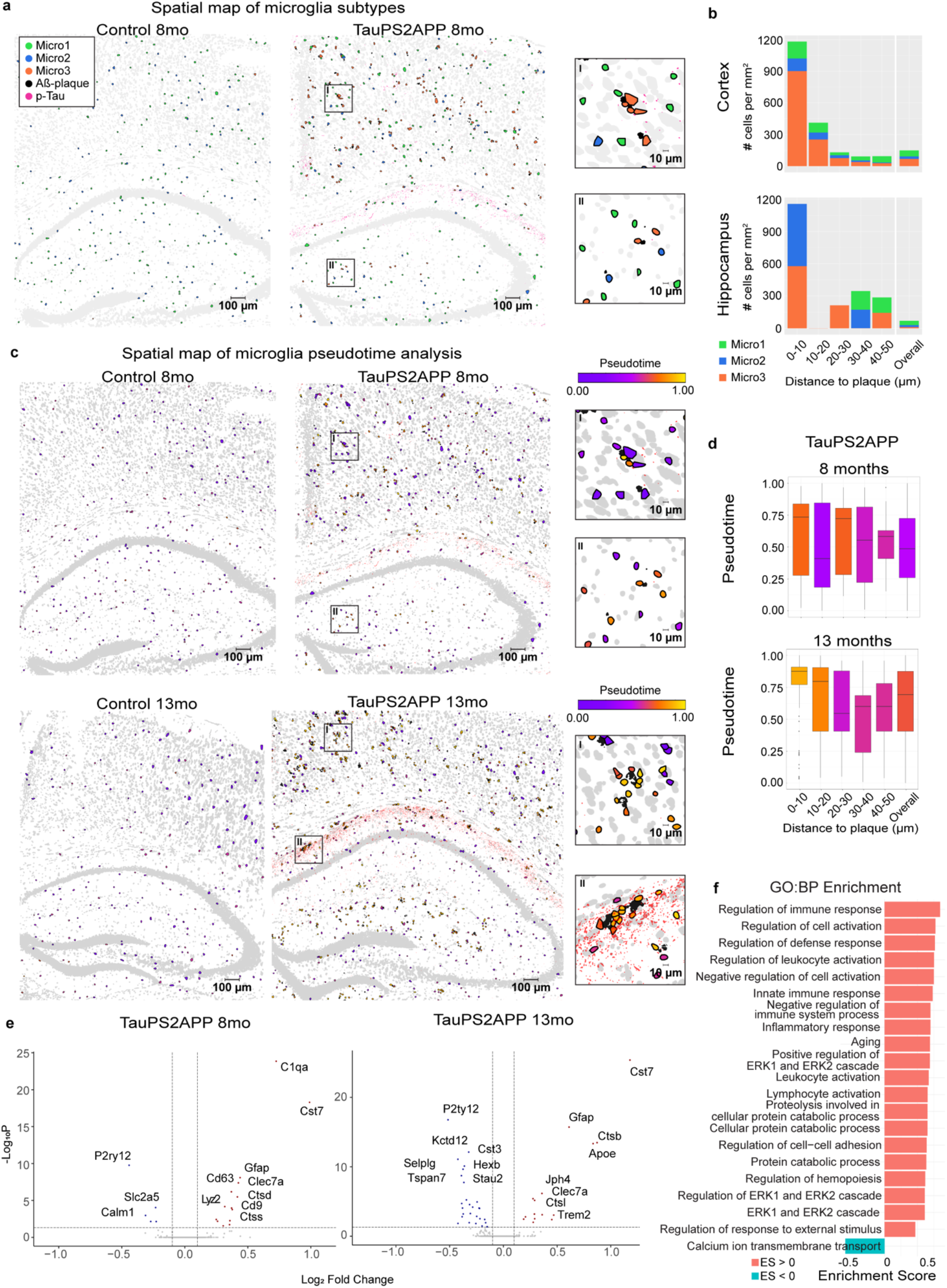
Additional gene expression and spatial analysis of microglia. **a,** Spatial map of microglia subtypes in control and TauPS2APP samples at 8 months. Scale bar, 100 µm. Two magnification sections (I, II) on the right side. Scale bar, 10 µm. **b,** Cell-type composition around Aβ plaque in different distance intervals for the TauPS2APP sample at 8 months. Stacked bar plot showing the density (count per mm^2^) of each microglia sub population in each distance interval (0-10, 10-20, 20-30, 30-40, 40-50 µm) around the Aβ plaque. The overall cell density of each subpopulation in each region was included as the reference for comparison. **c,** Spatial map of microglia colored by pseudotime. Scale bar, 100 µm. Two magnification sections (I, II) on the right side. Scale bar, 10 µm. **d,** Box plots showing the distribution of the pseudotime values of microglia in each distance interval (0-10, 10-20, 20-30, 30-40, 40-50 µm) around the Aβ plaque. A distribution of all the microglia was included as a reference. Box colors represent median pseudotime**. e,** Volcano plots for microglia differential expression. Plots showing the gene expression of microglia across AD and control samples in 8 and 13 months (y-axis: -log adjusted p-value, x-axis: average log fold change). Differentially expressed genes (p-value < 0.05, absolute value of logFC > 0.1) are marked in red (up-regulated) or blue (down-regulated). **f,** Gene set enrichment analysis (GSEA) result of differentially expressed genes (DEGs). Colored by sign of statistical significance (Nominal p-value < 0.01) enrichment score, Terms are filtered by term size: 20-1000.

**Extended Data Fig. 4.**
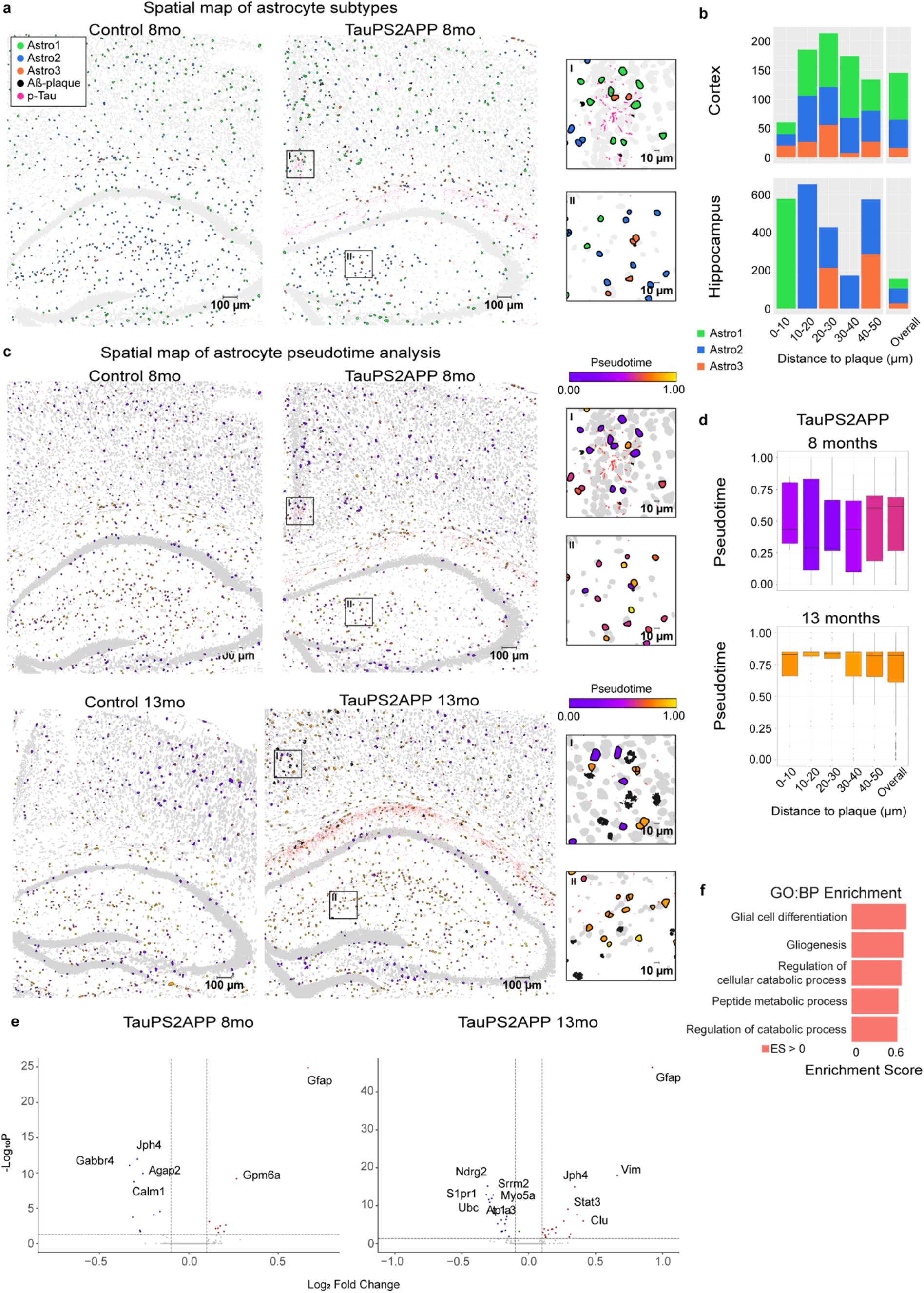
Additional gene expression and spatial analysis of astrocytes. **a,** Spatial map of astrocyte subtypes in Control and TauPS2APP samples in 8 month. Scale bar, 100 µm. Two magnification sections (I, II) on the right side. Scale bar, 10 µm. **b,** Cell type composition around Aβ plaque in different distance intervals for the TauPS2APP sample at 8 months. Stacked bar plot showing the density (count per mm^2^) of each astrocyte sub population in each distance interval (0-10, 10-20, 20-30, 30-40, 40-50 µm) around the Aβ plaque. The overall cell density of each subpopulation in each region was included as the reference for comparison (Overall).**c,** Spatial map of astrocytes colored by pseudotime for astrocyte population. Scale bar, 100 µm. Two magnification sections (I, II) on the right side. Scale bar, 10 µm. **d,** Box plots showing the distribution of the pseudotime values of astrocytes in each distance interval (0-10, 10-20, 20-30, 30-40, 40-50 µm) around the Aβ plaque. A distribution of all the astrocytes was included as a reference. **e,** Volcano plots for astrocytes differential expression. Plots showing the gene expression of astrocytes across AD and control samples in 8 and 13 months (y-axis: -log adjusted p-value, x-axis: average log fold change). Differentially expressed genes (p-value < 0.05, absolute value of logFC > 0.1) are marked in red (up-regulated) or blue (down-regulated). **f,** Gene set enrichment analysis (GSEA) result of differentially expressed genes (DEGs). Colored by sign of statistical significance (Nominal p-value < 0.01) enrichment score, Terms are filtered by term size: 20-1000.

**Extended Data Fig. 5.**
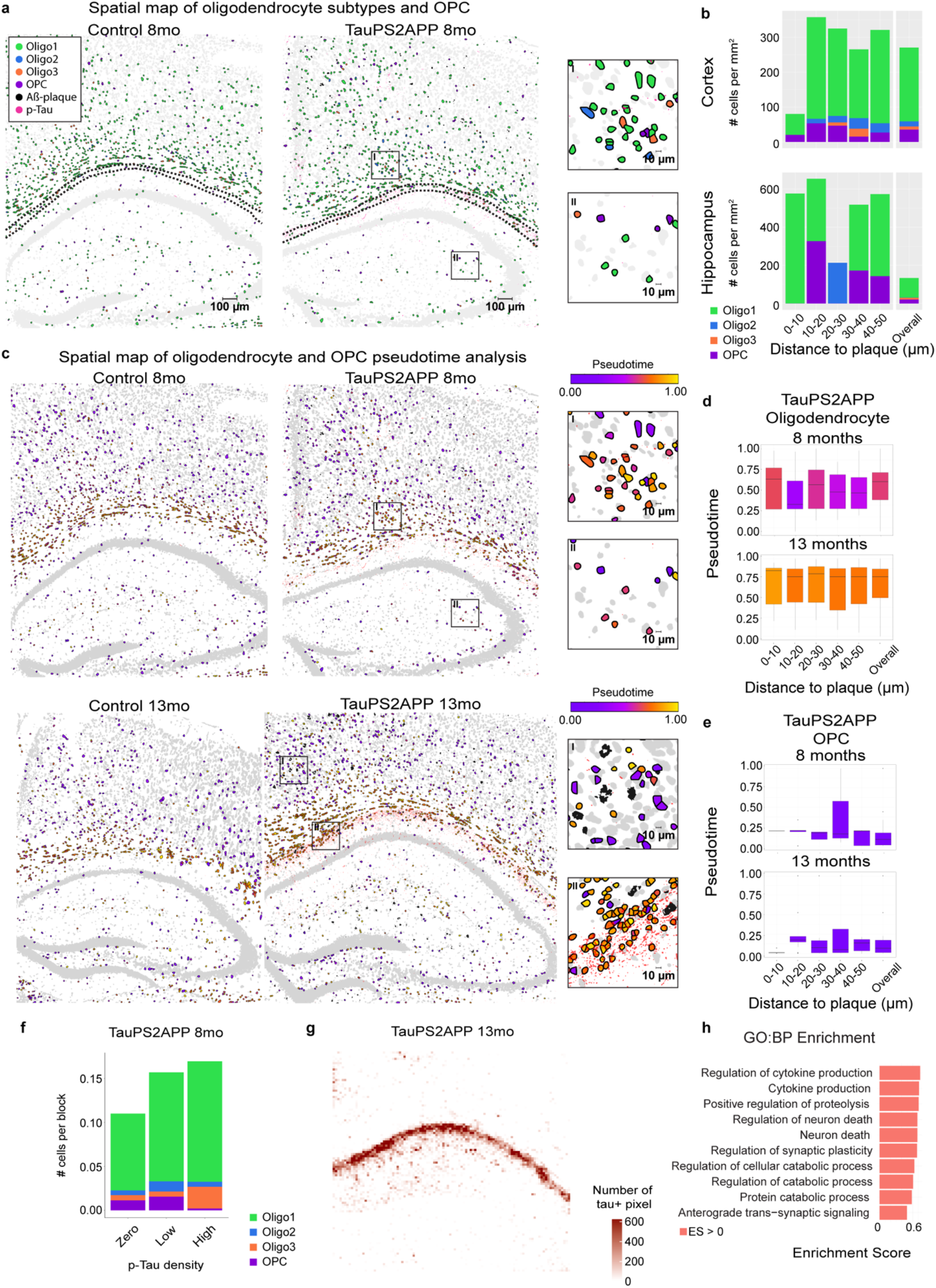
Additional gene expression and spatial analysis of oligodendrocytes and OPC. **a,** Cell-resolved spatial map for the oligodendrocyte and OPC population of Control and TauPS2APP mice at 8 months. Scale bar, 100 µm. Two magnification sections (I, II) on the right side. Scale bar, 10 µm. **b,** Cell type composition around Aβ plaque in different distance intervals for the TauPS2APP sample at 8 months. Stacked bar plot showing the density (count per mm^2^) of each oligodendrocyte sub-population and OPC in each distance interval (0-10, 10-20, 20-30, 30-40, 40-50 µm) around the Aβ plaque. The overall cell density of each subpopulation in each region was included as the reference for comparison (Overall).**c,** Spatial map colored by pseudotime for oligodendrocyte related population. Scale bar, 100 µm. Two magnification sections (I, II) on the right side. Scale bar, 10 µm. **d,** Pseudotime for oligodendrocytes in relation to plaque. Box plots showing the distribution of the pseudotime for oligodendrocytes in each distance interval (0-10, 10-20, 20-30, 30-40, 40-50 µm) around the Aβ plaque. A distribution of all oligodendrocytes was included as a reference. **e,** Pseudotime for OPC in relation to plaque. Box plots showing the distribution of the pseudotime for OPCs in each distance interval (0-10, 10-20, 20-30, 30-40, 40-50 µm) around the Aβ plaque. A distribution of all OPCs was included as a reference. **f,** Cell compositions in grid regions of the TauPS2APP sample at 8 month. Grid regions were ranked by the percentage of Tau positive pixels and fall into three bins: zero (0%), low (50%), high (100%). Stacked bar plot showing the average number of each sub-cluster of oligodendrocytes. **g,** Heatmap showing p-Tau quantification of each grid in TauPS2APP sample at 13 months. **h,** Gene set enrichment analysis (GSEA) result of differentially expressed genes (DEGs). Colored by sign of statistical significance (Nominal p-value < 0.01) enrichment score, Terms are filtered by term size: 20-1000.

**Extended Data Fig. 6.**
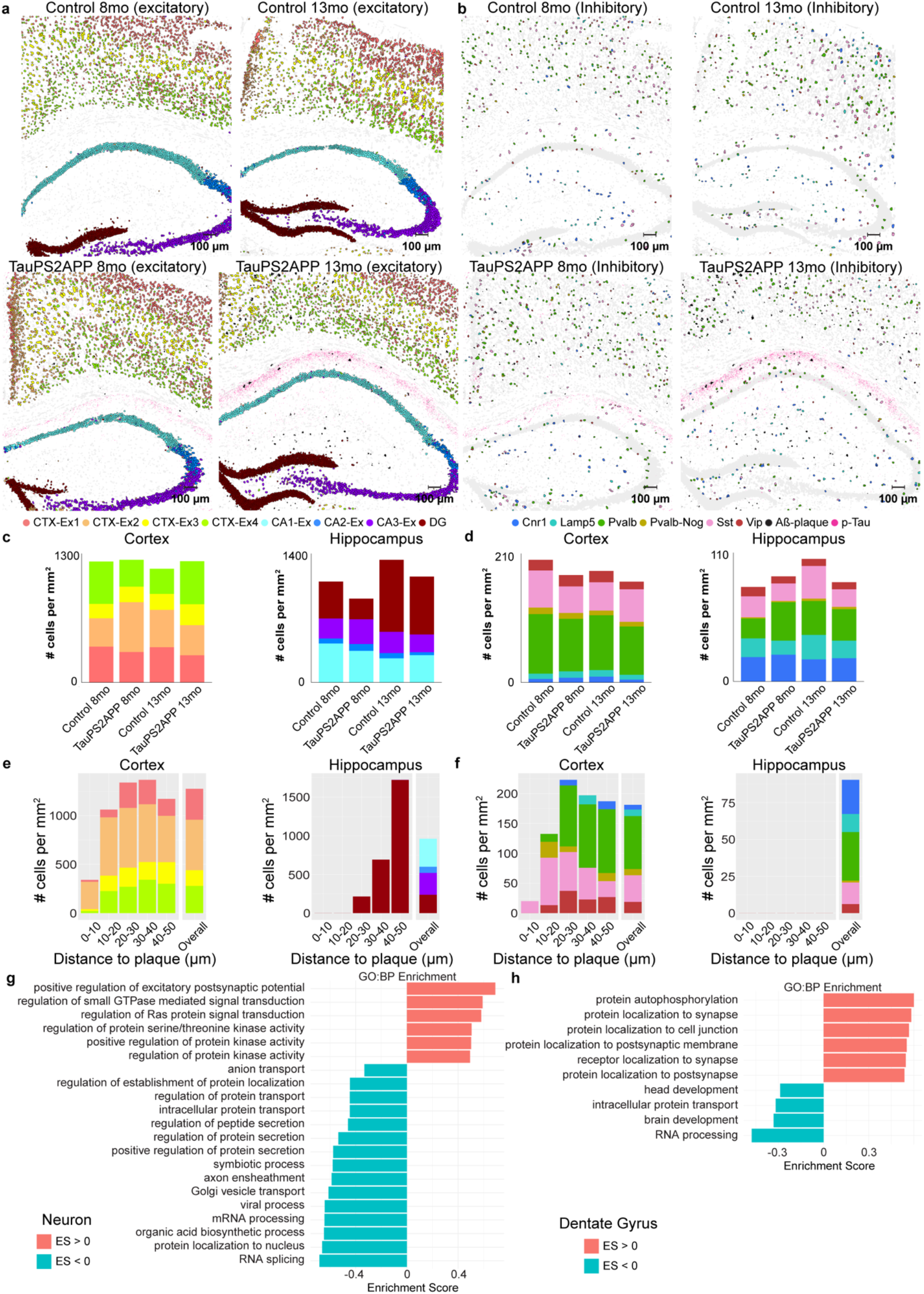
Additional gene expression and spatial analysis of neurons. **a,b,** Spatial maps for excitatory (**a**) and inhibitory (**b**) neuron populations of the 4 samples. Scale bar, 100 µm. **c,d,** Neuron density in each region. Stacked bar charts showing the density (count per mm^2^) of each sub-cluster of excitatory (**c**) and inhibitory (**d**) neuron population from the Cortex and Hippocampus region of all four samples. **e,f,** Neuron composition around plaque. Stacked bar charts showing the density (count per mm^2^) of each sub-cluster of excitatory (**e**) and inhibitory (**f**) neuron population from different brain regions at each different distance intervals (0-10, 10-20, 20-30, 30-40, 40-50 µm) to the Aβ plaque from the Cortex and Hippocampus region of TauPS2APP 8-month sample. The cell density of each subpopulation in each area was included as the reference for comparison (Overall). **g,** Gene set enrichment analysis (GSEA) results from differentially expressed genes (DEGs) in neurons. Colored by sign of statistical significance (Nominal p-value < 0.01) enrichment score, Terms are filtered by term size: 20-1000. **h,** Gene set enrichment analysis (GSEA) result of differentially expressed genes (DEGs) in neurons from Dentate Gyrus region. Colored by sign of statistical significance (Nominal p-value < 0.01) enrichment score, Terms are filtered by term size: 20-1000.

**Extended Data Fig. 7.**
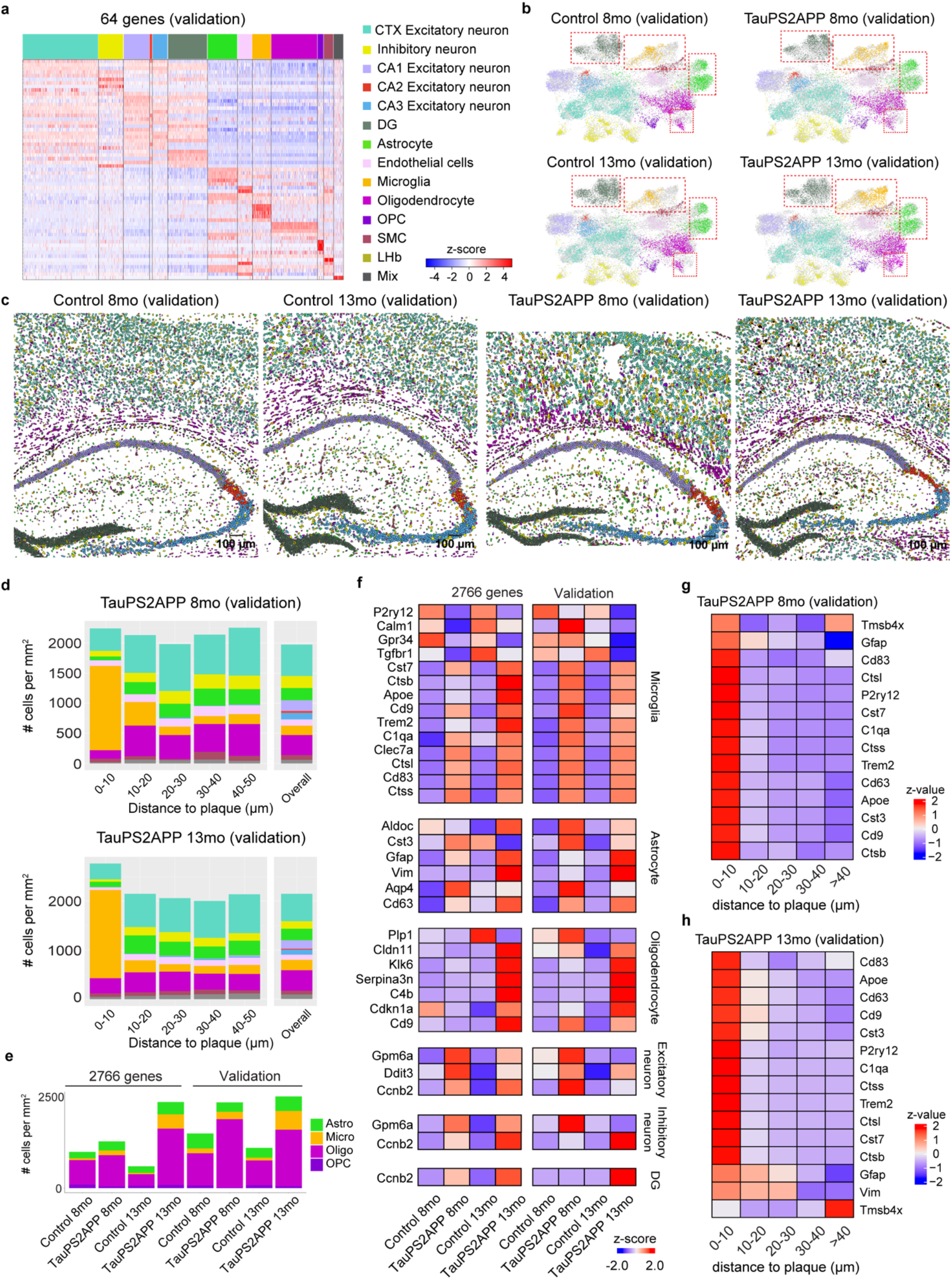
Validation of cell-type composition and spatial gene expression in TauPS2APP mice. **a,** Gene expression heatmaps for representative markers aligned with each top-level cell type of 64-gene (validation) datasets. Expression for each gene is z-scored across all genes in each cell. **b,** Uniform Manifold Approximation and Projection (UMAP) plot visualizing a non-linear dimensionality reduction for the transcriptomic profiles of 36,625 cells from 4 samples of the validation dataset. Plot showing the 13 top-level clusters: Cortex excitatory neuron (CTX-Ex, 8,640 cells) Inhibitory neuron (In, 2,858 cells), CA1 excitatory neuron (CA1-Ex, 2,967 cells), CA2 excitatory neuron (CA2-Ex, 331 cells), CA3 excitatory neuron (CA3-Ex, 1751 cells), Dentate Gyrus (DG, 4,560 cells), Astrocyte (Astro, 3,423 cells), Endothelial cell (Endo, 1,661 cells), Microglia (Micro, 2,183 cells), Oligodendrocyte (Oligo, 5,268 cells), Oligodendrocyte precursor cell (OPC, 711 cells), Smooth muscle cell (SMC, 1,146 cells), Mixed unidentified cells (Mix, 1,126 cells). **c,** Spatial atlas of top-level cell types in cortex and hippocampus regions of 4 samples in the 64-gene dataset. Scale bars, 100 µm. **d,** Cell-type composition around Aβ plaque at different distance intervals in both 8- and 13-month samples of the 64-gene validation dataset. Stacked bar plot showing the density (count per mm^2^) of each top-level cell type in each distance interval (0-10, 10-20, 20-30, 30-40, 40-50 µm) around the Aβ plaque. The overall cell density of each cell type in each region was included as the reference for (Overall). **e,** Barplot showing the cell density of astrocyte, microglia, oligodendrocyte and OPC in hippocampus alveus region in the 2,766- gene samples and validation samples. **f,** Matrix plot showing the row-wise scaled expression values of top significantly altered (rank by *p*-value) DEGs of glial cells and neuronal cells from TauPS2APP versus control samples**. g,h,** Matrix plot showing Plaque-Induced Genes (PIG, enriched in the 0-40 µm region, adjusted *p*-value < 0.01) that overlapped with 64-gene in each distance interval (0-10, 10-20, 20-30, 30-40 µm) around the Aβ plaque in the 8-month (H) and TauPS2APP 13-month sample (I) of the 64-gene validation dataset. Colored by row-wise *z*-score.

